# *In vivo* cortical neuron-astroglial functional coupling strengthens with acute stress but is impaired by chronic stress in mice

**DOI:** 10.1101/2025.09.07.674758

**Authors:** Yashika Bansal, Makayla A Mitchell, Sierra A Codeluppi-Arrowsmith, Jaime K Knoch, Kevan P Clifford, Yuliya S Nikolova, Etienne Sibille, Mounira Banasr

## Abstract

Exposure to acute, repeated, or chronic stress elicits a spectrum of cellular changes ranging from adaptive to maladaptive, including in the prefrontal cortex (PFC) where both neurons and astroglia undergo morphological, cellular and molecular remodeling. However, whether these alterations translate into altered cell activity and neuron-astroglia communication and how such changes evolve with recurrent stress or chronic stress exposure remains poorly understood. To address these questions, we used dual-color *in vivo* fiber photometry to longitudinally record PFC neuronal and astroglial calcium (Ca²⁺) signals in the same mice during a brief (tail pinch), sustained (immobilization) acute stress, repeated homotypic stress across weeks, and unpredictable chronic mild stress (UCMS), tracking of same cellular network across time. We found that acute stressors elicited coordinated increases in neuronal and astroglial Ca²⁺ activity and enhanced their Ca^2+^ signal coupling. Repeated intermittent stress induced comparable per-cell-type responses but progressively strengthened coupling, suggesting adaptation. We also demonstrated that this trajectory is reversed under UCMS; neuronal Ca²⁺ reactivity to stress challenge was sensitized whereas astroglial reactivity was blunted. This was associated with a weakened intercellular coupling (−3.5 fold) and astroglia became progressively hyporesponsive to neuronal drive. UCMS-induced neuron-astroglia functional coupling impairment coincided with the onset of anhedonia- and anxiety-like behavioral deficits. These findings position impaired neuron-astroglial functional coupling as a potential substrate of stress-induced cortical dysfunction that may distinguish adaptive from maladaptive stress responses, offering a mechanistically tractable target for intervention in stress-related psychiatric disease such as depression, where dysfunction in both cell types has been consistently reported.

## Introduction

Psychosocial stress is a leading risk factor for the onset and trajectory of major depressive disorder (depression) and other psychiatric disorders [1], yet the biological responses to stress span the continuum from adaptation to pathology [2, 3]. Brief or intermittent stress, with adequate recovery, engages physiological and neural mechanisms that drive adaptive plasticity, restore homeostasis and improve responses to future stressors [4]. However, when stress becomes prolonged, intense, or unpredictable, this adaptive response can evolve into a maladaptive one that is associated with enduring reorganization of the neural circuits governing mood and emotion [2, 5–7]. Through this transition, chronic stress confers vulnerability to anxiety [8], and depression [1], among the most prevalent and disabling psychiatric illnesses worldwide [9].

Within the circuits and brain regions most sensitive to stress, the prefrontal cortex (PFC) plays a central role in shaping mood- and cognition-related behaviors through its top-down control over the cortico-limbic circuit [10] Preclinical studies consistently show that the PFC undergoes profound cytoarchitectural remodeling under chronic stress, affecting both neurons and astroglia including neuronal atrophy with loss of dendritic spines and synapses [11–14] and reduced density and altered expression of cell-specific proteins of astroglia, including glial fibrillary acidic protein (GFAP) [15–17]. Together, these cellular changes are thought to underlie the volumetric loss and disrupted functional connectivity reported in both chronic stress rodent models [18, 19] and clinical studies of depression [14, 20–22]. Beyond recapitulating these cellular changes, chronic stress models elicit anxiety- and depressive-like behaviors, including anhedonia in rodents [19, 21, 23]. Across stress models, unpredictable chronic mild stress (UCMS) is a leading paradigm for dissecting chronic stress-induced changes at the cellular and circuit levels, in order to advance our understanding of stress-related pathology [20, 24, 25].

Both acute and chronic stress engage PFC neurons and astroglia, evidenced by immediate-early gene expression changes [26–28] and electrophysiology and other cell activity recording methods [29–31]. PFC neurons classically show increased activity following acute stress [27, 32] and persistent hyperactivity following UCMS [26]. Astroglial responses, although less studied, also show increased activity after acute stress [28, 33] but reduced activity in chronic corticosterone model [31]. However, whether and how these responses involve coordination between neurons and astroglia, and how repeated exposures progressively alters these responses remains poorly characterized.

Causal studies have independently implicated both populations in chronic-stress-related behavioural deficits. Activation or inhibition of specific PFC neuronal populations modulates anxiety- and depressive-like behavior [34], while astroglial loss or functional impairment is sufficient to induce chronic stress-like deficits [17, 35, 36]. Yet these cells do not act in isolation; cortical information computation depends on tight, bidirectional dialogue in which neurons propagate rapid signals across the synaptic cleft and neighboring astrocytes actively maintain synaptic homeostasis, respond through gliotransmission and provide metabolic support [17, 37, 38]. Calcium (Ca²⁺) dynamics are critical to this exchange as neuronal Ca²⁺ influx orchestrates synchronized neurotransmitter release, and astroglial Ca²⁺ transients drive gliotransmitter mobilization that modulate local synaptic plasticity [39, 40]. Temporal dynamic monitoring of Ca²⁺ activity therefore provides a high-fidelity readout of functional coordination within the neuron-astroglia network [41].

*In vivo* fiber photometry allows Ca^2+^ transients to be captured within molecularly defined cell types, yielding insights into the temporal dynamics of neuronal or astroglial cell activity associated with behavior and experimental manipulations [42–46]. Advances in dual- and multi-color fiber photometry have extended this capability and now both cell types and their potential coordinated dynamic responses are being explored revealing that neuronal and astroglial Ca²⁺ signals are tightly coupled, synchronized or phase-shifted depending on brain regions or behavior examined [43, 47]. However, how this dynamic coupling is affected by acute, repeated or chronic stress remain largely unknown, leaving a key cellular dimension of stress neurobiology unresolved, especially whether coordinated neuron-astroglia dynamics can provide insights into adaptive and maladaptive stages associated with stress-related vulnerability.

Here, we investigated neuronal and astroglial Ca²⁺ dynamics as well as their functional coupling across a graded series of stress exposures differing in nature, duration, recurrence, and chronicity, with the goal of characterizing the cell activity signatures within the PFC neuron-astroglia network distinguishing adaptive from maladaptive stress responses. First, we examined how acute stress (brief or sustained) alters neuronal and astroglial Ca^2+^ activity in the PFC, recorded separately and concurrently; simultaneous recording of both cell populations further enabled us to assess cooperative Ca²⁺ dynamics as an index of functional coupling. Second, we characterized how repeated acute stress affects individual cell-type signaling, coupling strength, and the directionality of information flow between neurons and astroglia, probing the intercellular mechanisms engaged during adaptation to recurrent intermittent homotypic stress challenges. Finally, using the UCMS model, we assessed the longitudinal consequences of chronic stress on baseline neuronal and astroglial Ca^2+^ activity, and their coupling as well as the PFC neuron-astroglia network’s response to weekly re-exposure to acute stress challenges. In parallel, anhedonia and anxiety-like behaviors were monitored enabling us to map cell activity changes onto the emergence of UCMS behavioral deficits. We found that whereas acute stress enhances neuron-astroglia functional coupling as part of an adaptive response, chronic stress disrupts this bidirectional communication, inducing a breakdown within the network that either precedes or emerges alongside depressive-like deficits.

## Results

### Acute tail pinch stress increases PFC neuronal and astroglial Ca^2+^ activity

To confirm prior single-channel fiber photometry studies reporting increases of activity in neurons [48] or astroglia [49] following tail pinch, we first investigated *in vivo* modulation of PFC neuronal or astroglial Ca^2+^ signals individually during acute stress (Figure S1) using AAVs driving the expression of the genetically encoded Ca^2+^ indicator GCaMP6f under either the neuron-specific human synapsin1 (Syn1) promoter (AAV-Syn1-GCaMP) or the GFAP+ astroglia-specific GfaABC1D promoter (AAV-GFAP-GCaMP) (n=8 per cohort). Acute tail pinch robustly triggered PFC neuronal or astroglial Ca²⁺ transients (supplementary results and Figure S1).

To specifically investigate the temporal relationship between neuronal and astroglial responses, we next examined Ca^2+^ transients in both cell types simultaneously using dual-color fiber photometry. Mice were co-injected with two viruses expressing either jRCaMP1a under the Syn1 promoter (AAV-Syn1-RCaMP) and GCaMP6f under the GfaABC1D promoter (AAV-GFAP-GCaMP); out of 16 mice surgerized, 14 mice yielded analyzable signals for both channels up to the tail-pinch challenge (Figure 1A). Viral specificity was confirmed by immunohistochemistry and infection rates were quantified (Figure 1D-H). Consistent with the single-color experiments, analysis of the magnitude and kinetics of Ca^2+^ signals across the three windows: pre-pinch (−5 to 0 s), pinch (0 to 3 s), and post-pinch (3 to 20 s) showed significant increases in tail pinch-evoked mean z-scored ΔF/F in both cell types. Specifically, repeated-measures one-way ANOVA revealed a significant main effect of time window on neuronal Ca^2+^ signals (F_(2,26)_=10.05, *p*=0.0006; Figure 2A, C and D) and astroglial Ca^2+^ signals F(_2,26)_=19.69, *p*<0.0001; Figure 2B, C and E) reflecting the significant increase in both cell types during tail pinch (neurons: p<0.001; astroglia: p<0.01 vs pre-pinch, Tukey’s *post-hoc* test) and post-pinch (neurons: p<0.01; astroglia: p<0.0001 vs pre-pinch). Parallel increases were found on integrated signal i.e. area under the curve (AUC) (neurons: F_(2,26)_=47.78, *p*<0.0001; astroglia: F_(2,26)_=40.50, *p*<0.001) during and post-pinch relative to baseline (Figure 2D and E). Importantly the temporal patterns of Ca^2+^ transients differed between cell types: astroglial responses showed a slightly longer onset latency than neuronal responses (mean lag=0.32s; Figure 2C). Including sex as a factor in the analyses revealed no main effect of sex (neurons: F_(1,12)_=0.00; astroglia: F_(1,12)_=0.08) or sex x time interaction (neurons: F_(2,24)_=1.41; astroglia: F_(2,24)_=0.56), and confirmed the main effect of time (neurons: F_(2,24)_=15.09; astroglia: F_(2,24)_=11.09, *p*<0.001), reflecting increase in tail pinch-evoked Ca^2+^ signals in both cell types.

**Figure 1.**
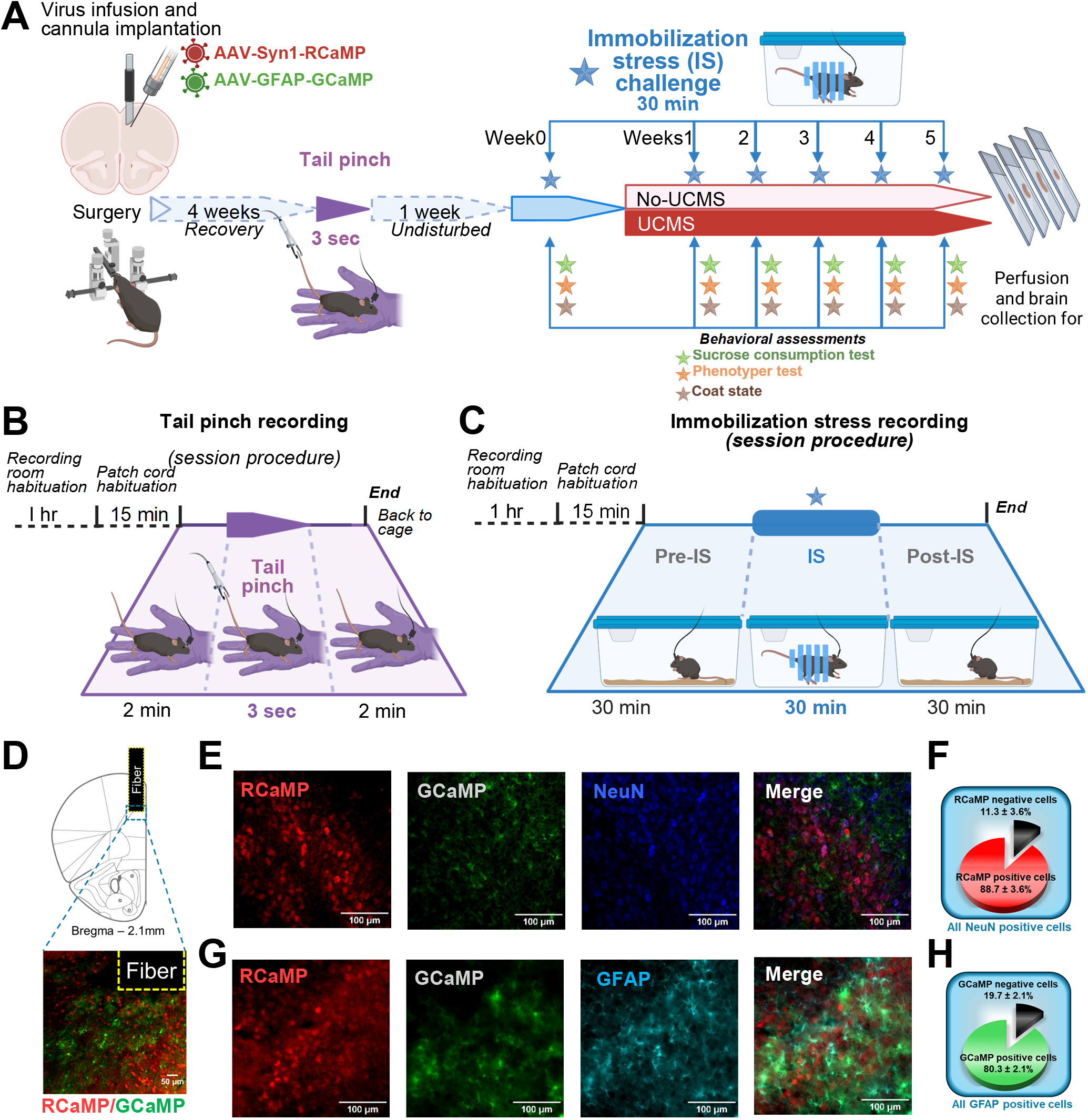
Experimental design, stress and recording procedures, cell specificity and infection rate of viruses. **(A)** Timeline from surgery to brain collection. **(B)** Tail-pinch session recording procedure. **(C)** IS session recording procedure. Panels A-C were created in https://BioRender.com. **(D)** Atlas schematic of recording sites (top) and representative image of cannula placement in the PFC (bottom). **(E-H)** Representative images of AAV-Syn1-RCaMP and AAV-GFAP-GCaMP infected areas co-labelled with either NeuN **(E)** or GFAP **(G)** With pie charts showing quantified infection rates for neurons (NeuN+/RCaMP+ vs NeuN+/RCaMP-,) **(F)** and astroglia (GFAP+/GCaMP+ vs GFAP+/GCaMP-) **(H)**.

**Figure 2.**
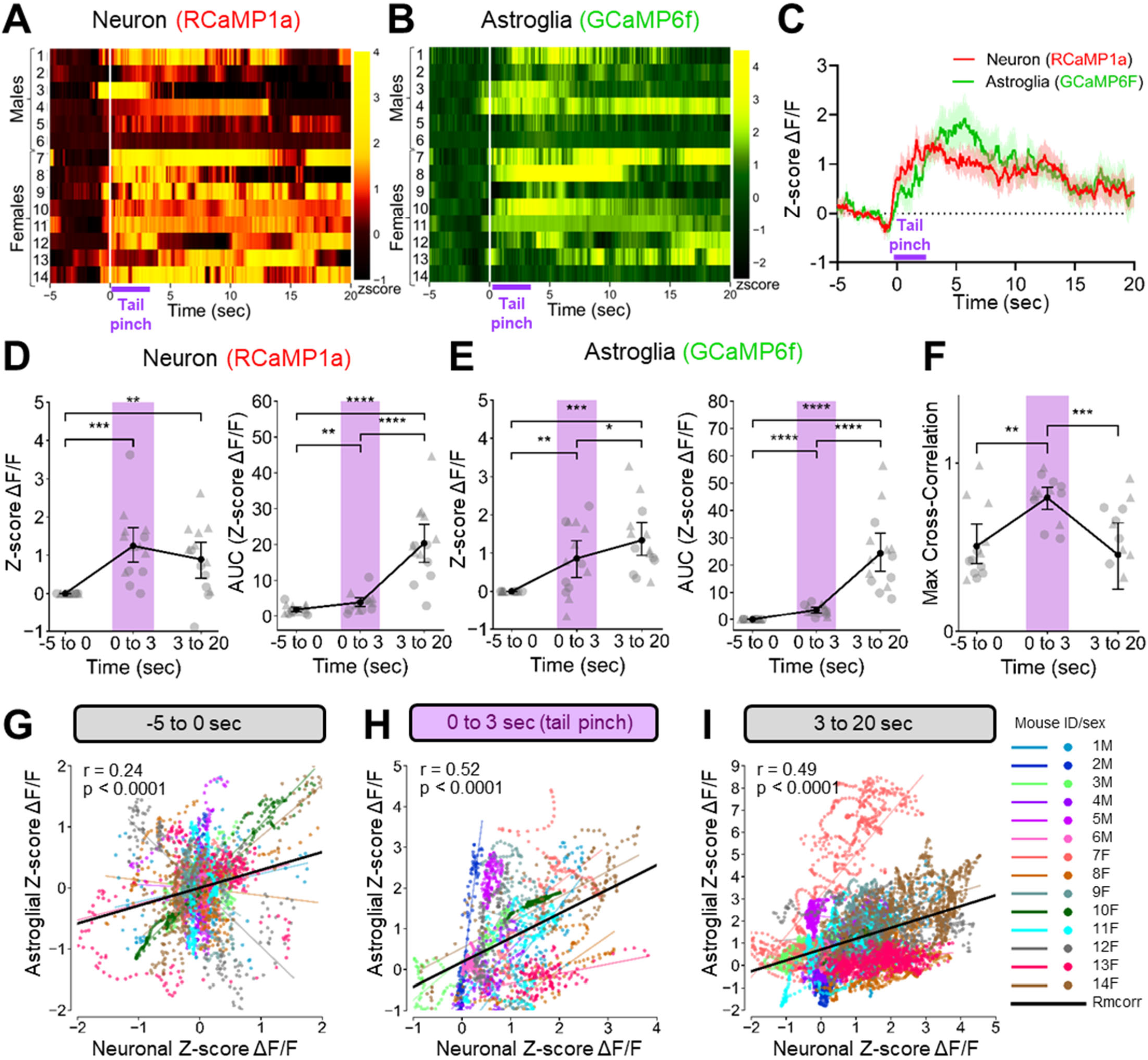
Tail pinch increases neuronal and astroglial Ca^2+^ activity in mouse PFC. (**A-B)** Heatmaps of zscore ΔF/F of neuronal **(A)** and astroglial **(B)** Ca^2+^ signals recorded for each mouse (row) with white line indicating the tail pinch onset and purple bar pinch duration. **(C)** Average zscore ΔF/F of Ca^2+^ signals from PFC neurons (red) and astroglia (green). Solid lines indicate mean and shaded areas indicate s.e.m. **(D-E)** Average zscore ΔF/F (left) and area under curves (AUC) (right) of neuronal **(D)** and astroglia Ca2+ signals **(E)** calculated for each time period. Individual data points represent single subjects (circles for males, triangles for females). Black dots and error bars indicate mean ± s.e.m. **(F)** Max cross-correlation between neuronal and astroglial Ca^2+^ signals. *p<0.05, **p<0.01; ***p< 0.001; ****p<0.0001. **(G-I)** Scatter plot of neuronal z-score ΔF/F and astroglial z-score ΔF/F (**G**: -5 to 0 sec; **H**: 0 to 3 sec, tail pinch; **I**: 3 to 20 sec). Each color represents individual data points (50 points per sec) for each mouse. Color lines represent the Pearson’s regression slope for each mouse. The bold black line represents the repeated measure correlation (*rmcorr*) indicating common within-subject association with r and p values reported in each panel.

To investigate whether acute stress alters functional coupling between neurons and astroglia, we quantified the temporal coordination of their Ca²⁺ signals using the maximum cross-correlation between concurrent z-scored ΔF/F traces within each time window (Figure 2F). Peak cross-correlation increased markedly during tail pinch relative to both pre- and post-pinch (F_(2,26)_=7.85, *p*=0.002), indicating that acute stress not only amplified the magnitude of activity in both cell types (Figure 2D–E) but also enhanced their temporal coordination.

Because cross-correlation reflects maximum coupling at the population level but does not isolate moment-to-moment coupling within individual animals, we complemented this analysis by applying repeated-measures correlation (*rmcorr*) [50]. *Rmcorr* accounts for repeated observations and quantifies within-subject coupling between two signals for each time window. This analysis revealed a significant within-subject coupling for all three conditions, demonstrating that fluctuations in neuronal Ca^2+^ signals were reliably tracked by corresponding astroglial Ca^2+^ changes (Figure 2G–I). The strength of this coupling doubled during tail pinch (r=0.52 vs. pre-pinch r=0.24) and remained elevated post-stimulus (r=0.49). There were no sex differences in neuron-astroglia functional coupling assessed using either maximum cross-correlation or *rmcorr*. Notably, the post-pinch divergence between the cross-correlation and *rmcorr* results suggest that acute stress produces two qualitatively distinct effects on neuron-astroglia coupling: a rapid, stimulus-locked coordination at the population level (cross-correlation), and a slower, sustained within-animal coupling that persists beyond the stressor (*rmcorr*). Together, the two analyses converge to indicate that acute stress induces a strong increase in functional coupling between PFC neurons and astroglia.

### Immobilization stress drives sustained increases in PFC neuronal and astroglial Ca²⁺ activity

We next assessed the effects of a 30 min sustained immobilization stress (IS) on PFC neuronal and astroglial Ca²⁺ signals, recorded both individually and concurrently. Each recording session comprised three continuous 30 min phases: pre-IS, IS, and post-IS conditions (Figure 1A, C). Cohort sizes were smaller between the tail pinch and IS experiments due to loss of cannula or signal quality, yielding n=7 for Syn1-GCaMP, n=8 for GfaABC1D-GCaMP cohorts (Figure S2), and n=12 for the combined Syn1-jRCaMP1a + GfaABC1D-GCaMP cohort (Figure 3). IS-evoked response obtained with single-channel recordings were similar to dual-channel recordings and are presented in supplementary material and Figure S3.

**Figure 3.**
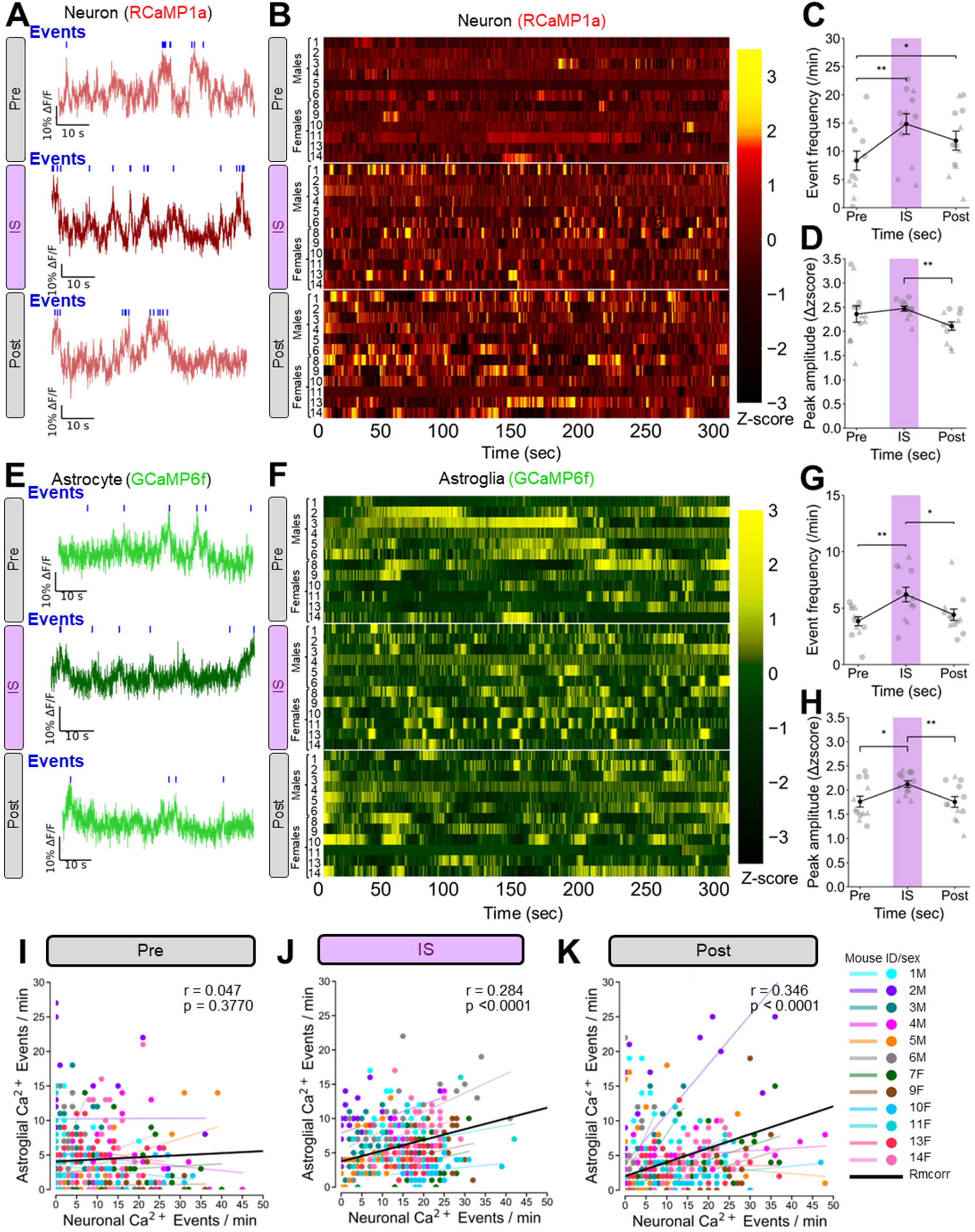
Immobilization stress (IS) challenge increases neuronal and astroglial Ca^2+^ activity in mouse PFC. (A, **E)** Representative traces of simultaneously recorded neuronal **(A)** and astroglial **(E)** Ca^2+^ signals (1 min-duration) obtained from one mouse for both channels during Pre-IS, IS and Post-IS conditions. Blue ticks represent the above threshold detected Ca^2+^ events. (**B, F)** Heatmaps of zscore ΔF/F (5 min-duration) of neuronal **(B)** and astroglial Ca^2+^ **(F)** signals obtained for each mouse (row). Chosen 5 min windows are the last 5min of pre-IS and the 5 first minutes of IS and Post-IS challenge. **(C, G)** Ca^2+^ event frequency for neurons **(C)** and astroglia **(G)** during three conditions of IS challenge. **(D, H)** Event peaks amplitude neurons **(D)** and astroglia **(H)**. Individual data points represent single mouse (circles for males, triangles for females). Black dots and error bars indicate mean ± s.e.m. *p<0.05, **p<0.01 **(I-K)** Scatter plot of neuronal Ca^2+^ events and astroglial Ca^2+^ events during the three conditions Pre **(I)**, IS **(J)** and Post **(K).** Each color represents individual data points (1 per minute) for each mouse. Color lines represent the Pearson’s regression slope for each mouse. The bold black line represents the repeated measure correlation (*rmcorr*) indicating common within-subject association with r and p values reported in each panel.

Concurrent dual-color recording in the Syn1-jRCaMP1a + GfaABC1D-GCaMP cohort (Figure 3A-B and E-F) showed that IS triggered a significant increase in neuronal and astroglial Ca^2+^ event frequency. Indeed, repeated measures ANOVA showed a main effect of time in both cell types (neurons: F_(2,22)_=12.44, *p*=0.0008; astroglia: F_(2,22)_=10.02, *p*=0.0008). Neuronal Ca^2+^ event frequency significantly increased during IS (∼65%, *p*<0.01 vs Pre-IS; Tukey’s *post-hoc* test) and remained elevated post-IS (*p*<0.05; Figure 3C). Astroglial Ca^2+^ event frequency significantly increased (∼50%, *p*<0.01 vs pre-IS) and returned to baseline during post-IS condition (*p*<0.05) (Figure 3G). Similar effects were found on event amplitude (neurons: F_(2,22)_=4.73, *p*=0.031, Figure 3D; astroglia: F_(2,22)_=4.50, *p*=0.032, Figure 3H). Including sex as a factor in the analyses showed no main effect of sex (neurons: F_(1,10)_=0.86; astroglia: F_(1,10)_=0.16) or sex x time interaction (neurons: F_(2,20)_=0.50; astroglia: F_(2,20)_=0.46), and confirmed the main effect of time (neurons: F_(2,20)_=7.47; astroglia: F_(2,20)_=9.02, *p*<0.01) reflecting increase in Ca^2+^ events in both cell types during IS.

Notably, the pattern of astroglial Ca²⁺ activity differed markedly between baseline and IS (Figure 3F). Pre-IS astroglial signals were dominated by prolonged periods of sustained activity alternating with extended quieter periods; during IS, this pattern shifted toward discrete, short-duration bursts of Ca^2+^ activity that seem to map onto neuronal activity bursts (Figure 3B). These astroglial activity changes in activity pattern were overall superimposed onto a noisier background than that of neurons (Figure 3A and E). To address potential changes in coordination between neuronal and astroglial Ca^2+^ signals, we next examined how IS affects neuron-astroglia functional coupling. To do so, we applied *rmcorr* to event frequency per minute (Figure 3I–K). This approach is better suited to 30-min recordings than cross-correlation on continuous ΔF/F, which over long windows is dominated by slow shared trends in activity; *rmcorr* instead isolates within-animal coupling from paired fluctuations in event rate. We found that neuronal and astroglial Ca²⁺ event rates over 30 min window showed no significant association at pre-IS (r=0.05, *p*=0.38), became significantly correlated during IS (r=0.28, *p*<0.0001), with the correlation strengthening further in the post-IS condition (r=0.35, *p*<0.0001). Thus, IS increased neuron-astroglia functional coupling both during and after sustained stress exposure.

### Chronic stress exposure differentially alters PFC neuronal and astroglial Ca^2+^ responses to IS challenge

A central question in stress neurobiology is how chronic stress alters not just baseline activity but the dynamic response of cortical circuits to subsequent stressful life events. Addressing this question requires tracking the same cells or cell population across the full trajectory of chronic stress - an approach unprecedently applied to investigate neuron-astroglia interactions and functional coupling. Here we recorded neuronal and astroglial Ca²⁺ activity from the same animals each week across 5 weeks of unpredictable chronic mild stress (UCMS, n=6) and compared to no-stress conditions (no-UCMS, n=6) (Figure 1A). Each weekly recording session consists of pre-IS, IS, and post-IS periods (30 min each) allowing us to track, week by week, how prior chronic stress reshapes the previously-characterized acute stress response of each cell type and the functional coupling between them.

As our UCMS protocol [51–53] had to be adapted for fiber-optic-implanted animals, we monitored physical changes (coat state), anhedonia-like behavior (sucrose consumption test) and anxiety-like behavior (Phenotyper test) longitudinally throughout the chronic stress exposure. Behavioral analysis confirmed that UCMS induced depressive-like behaviors by 4-5 weeks (Figure 4A-C). UCMS progressively degraded coat state across weeks (stress x time interaction: F_(5,50)_=4.18, *p*<0.01, Figure 4A), reaching significance by week 5 (*p*<0.05, Tukey’s *post hoc*). Sucrose intake in the sucrose consumption test was reduced in UCMS mice but only at week 5 (*t*-test, *p*<0.05), with no accompanying change in water consumption (Figure S3); the corresponding change in sucrose preference was small and non-significant (data not shown). Using the PhenoTyper test, which allowed to measure anxiety-like response to light challenge applied weekly, we also assessed post-light challenge avoidance behavior i.e. residual avoidance as per Prevot et al [16, 54] (Figure S3); UCMS progressively increased residual avoidance of the lit zone (stress × time interaction: F_(5,50)_=12.87, *p*<0.05; Figure 4C), reaching significance at week 4 (*p*<0.001) and week 5 (*p*<0.01, Tukey’s *post hoc*). Together, these changes confirm that the modified protocol reproduced the hallmark anhedonia, anxiety-like behaviour, and physical deterioration characteristic of chronic stress paradigms [16, 35, 51, 55] despite the constraints of fiber-optic implantation and repeated longitudinal recording. Importantly, the same weekly IS recordings were applied to the no-UCMS group explaining the slight increase in coat state degradation in that group, yet robust differences between no-UCMS and UCMS animals were still detected across all of these behavioral measures.

**Figure 4.**
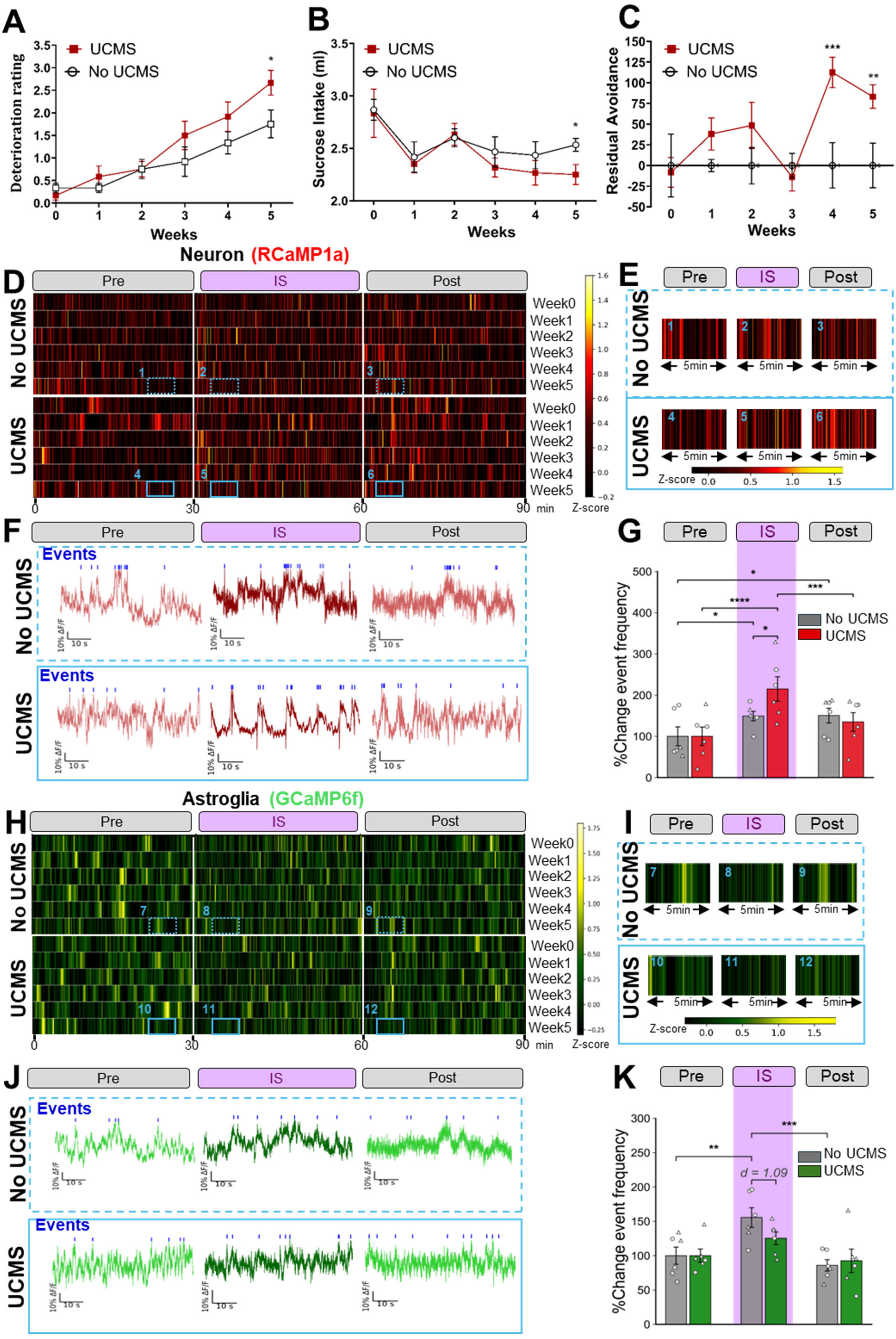
UCMS induced depressive-like behaviors, exacerbated PFC neuronal Ca^2+^ activity, and impaired astroglial activity in response to IS challenge. (A-C) Graphs representing the data obtained during the weekly coat state **(A)**, sucrose test **(B)** and PhenoTyper test **(C)**. Data are represented as mean ± s.e.m. **(D, H)** Heatmap of average zscore ΔF/F of neuronal **(D)** and astroglial **(H)** Ca^2+^ activity in no-UCMS and UCMS group during three conditions Pre-IS, IS, and Post-IS on week 0,1,2,3,4 and 5. **(E, I)** Magnified heatmap displaying a 5-minute representative window of neuronal **(E)** and astroglial **(I)** Ca^2+^ activity at week 5. **(F, J)** Representative traces of simultaneously recorded neuronal **(F)** and astroglial **(J)** Ca^2+^ signals (1 min-duration) obtained from one mouse for both channels during Pre-IS, IS and Post-IS conditions. Blue ticks represent the detected Ca^2+^ events. **(H, M)** Histograms representing % change in neuronal **(G)** and astroglial **(K)** event frequency normalized to pre-IS condition for each animal and for each group on Week 5. *p<0.05, **p<0.01, ***p<0.001, ****p<0.0001.

Throughout we examined how UCMS affects neuronal and astroglial Ca²⁺ signaling during the weekly IS challenges. In the no-UCMS control group, recurrent exposure to IS reliably increased neuronal Ca²⁺ event frequency and amplitude each week (1.4 – 2-fold change relative to pre-IS; Figure 4D and S4), with parallel but more modest increases in astroglial Ca^2+^ event rates (30-59% relative to pre-IS, Figure 4H and S5). Both no-UCMS and UCMS-exposed mice showed the same direction of response in both cell types through weeks 1–3, however, by weeks 4 (Figure S4 and S5) and 5 (Figure 4G and K) the two groups diverged in a cell-type-specific manner. More precisely, when analyzing percent change in peak frequency normalized to pre-IS condition, we found that IS-induced increase in neuronal Ca^2+^ event frequency was approximately 2-fold larger in UCMS animals compared to no-UCMS controls (Figure 4E–G; two-way ANOVA: stress × time interaction F_(2,20)_=6.50, *p*<0.01; UCMS vs no-UCMS during IS, *p*<0.05; Figure 4G) whereas the corresponding astroglial response was attenuated. Indeed, no-UCMS controls showed a robust ∼55% increase in astroglial event frequency during IS, while UCMS animals showed no significant IS response (Figure 4I–K). Importantly, although not statistically significant the difference between the blunted astroglial Ca^2+^ response to IS was biologically meaningful (during IS: UCMS vs No-UCMS Cohen’s *d*=1.09; Figure 4K). In sum, chronic stress drives opposing cell-type-specific changes in PFC Ca^2+^ signaling reactivity to acute stress challenge: neuronal sensitization and astroglial blunting.

To verify that these group differences were not attributable to disparities in viral expression, we compared infection rates and infected cell densities between groups. Neither infection rates (no-UCMS: 89.6 ± 5.6% vs. UCMS: 87.8 ± 5.3% for AAV-syn1-RCaMP; no-UCMS: 79.5 ± 2.8% vs. UCMS: 82.1 ± 7.7% for AAV-GFAP-GCaMP) nor densities of infected cells (no-UCMS: 95,732 ± 9,137 vs. UCMS: 100,545 ± 11,882 cells/mm³ for neurons; no-UCMS: 21,000 ± 600 cells/mm³ vs. UCMS: 23,100 ± 1,950 cells/mm³ for neurons) differed between groups, indicating that the sensitized neuronal and blunted astroglial Ca²⁺ responses were not attributable to altered viral expression or cell loss.

To determine whether these divergent cell-type responses also affected functional coupling, we examined the association between neuronal and astroglial Ca^2+^ event rates separately for each condition (Pre, IS, Post) and each week (W0–W5) using *rmcorr* (Table 1 and Figure S6). In no-UCMS control group, repeated IS exposure induced a progressive strengthening of neuron-astroglia coupling in pre-IS conditions, emerging from week 2 onward. During IS, the first exposure enhanced neuron-astroglia coupling, which disappeared on week 1 and 2 and restored on week 3 – 5 while post-IS coupling was robust at every week (Table 1A). Interestingly, UCMS-exposed animals showed a strikingly different pattern: Pre-IS baseline coupling was irregular in UCMS animals, IS-condition coupling was abolished from week 2 onward (Table 1B), and post-IS coupling remained significant at every week although attenuated in magnitude relative to no-UCMS control group. Here, across group comparisons are descriptive, but suggest that recurrent acute stress exposure progressively strengthens neuron-astroglia coupling across all three conditions (Pre, IS, Post), potentially reflecting an adaptive process that enhances communication between the two cell types. In contrast, prior chronic stress disrupts this adaptive strengthening, most strikingly during pre-IS and IS conditions and to a lesser extent post-IS. This phase-specific loss of acute stress-evoked coupling may indicate that UCMS disrupts the dynamic neuron-astroglia communication normally engaged in responding to acute stressful stimuli.

**Table 1.**
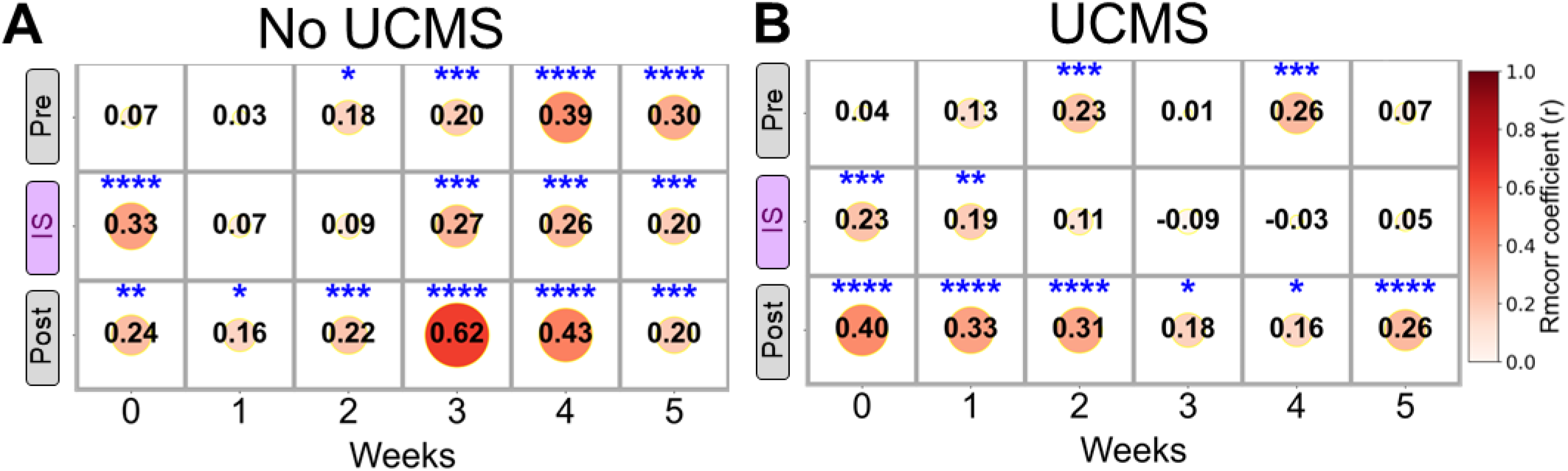
IS and UCMS alter neuronal and astroglial Ca^2+^ coupling strength during Pre, IS and Post-IS condition. r values obtained for each group and condition using *rmcorr,* *p<0.05, **p<0.01, ***p<0.001, ****p<0.0001.

### Neuronal Ca^2+^ activity precedes and drives astroglial activity during acute immobilization stress

Because *rmcorr* does not allow direct comparisons between groups or time points, our analyses up to this point characterized neuron-astroglia functional coupling without addressing two key dimensions: the direction of information flow between the two cell types, and how acute and chronic stress affect directional functional coupling within and between groups across time. We addressed both questions with complementary analytical approaches.

To establish the temporal directionality of the coupling between PFC neuronal and astroglial Ca^2+^ signals, we used three parallel methods: cross-correlation lag, Granger causality, and transfer entropy, in order to identify the dominant direction during first IS challenge at week0 (n=12; Figure 5A–D). Firstly, cross-correlation of decimated to 1 Hz z-scored ΔF/F signals revealed a clear directional asymmetry where neuronal activity preceded astroglia activity with a peak lag of +1 s and a mean per-animal cross-correlation peak of r=0.44 (Figure 5A-B), consistent across individual subjects rather than reflecting random co-variation. Secondly, Granger causality analysis, which uses linear autoregressive models to test whether one signal’s history predicts another’s future beyond what the target’s own history explains, yielded a significantly higher F-statistic for the Neu→Ast direction than for Ast→Neu (*p*<0.01; Figure 5C), confirming that neuronal Ca²⁺ activity carries greater predictive information about future astroglial activity than the reverse. Finally, because Ca²⁺ dynamics can be nonlinear and may be underestimated by linear methods, we further applied Transfer Entropy, a model-free measure of directional information flow that captures both linear and nonlinear dependencies. Transfer entropy corroborated this asymmetric coupling (asymmetry index = +43%), with Neu→Ast information flow 2.5-fold higher than the Ast→Neu (0.15 vs 0.06 bits), indicating predominantly neuron-led coupling (*p*<0.01; Figure 5D). Together, the convergence of cross-correlation lag, Granger causality, and transfer entropy provided three independent lines of evidence that, under IS challenge, neuronal activity drives astroglial activity in this circuit.

**Figure 5.**
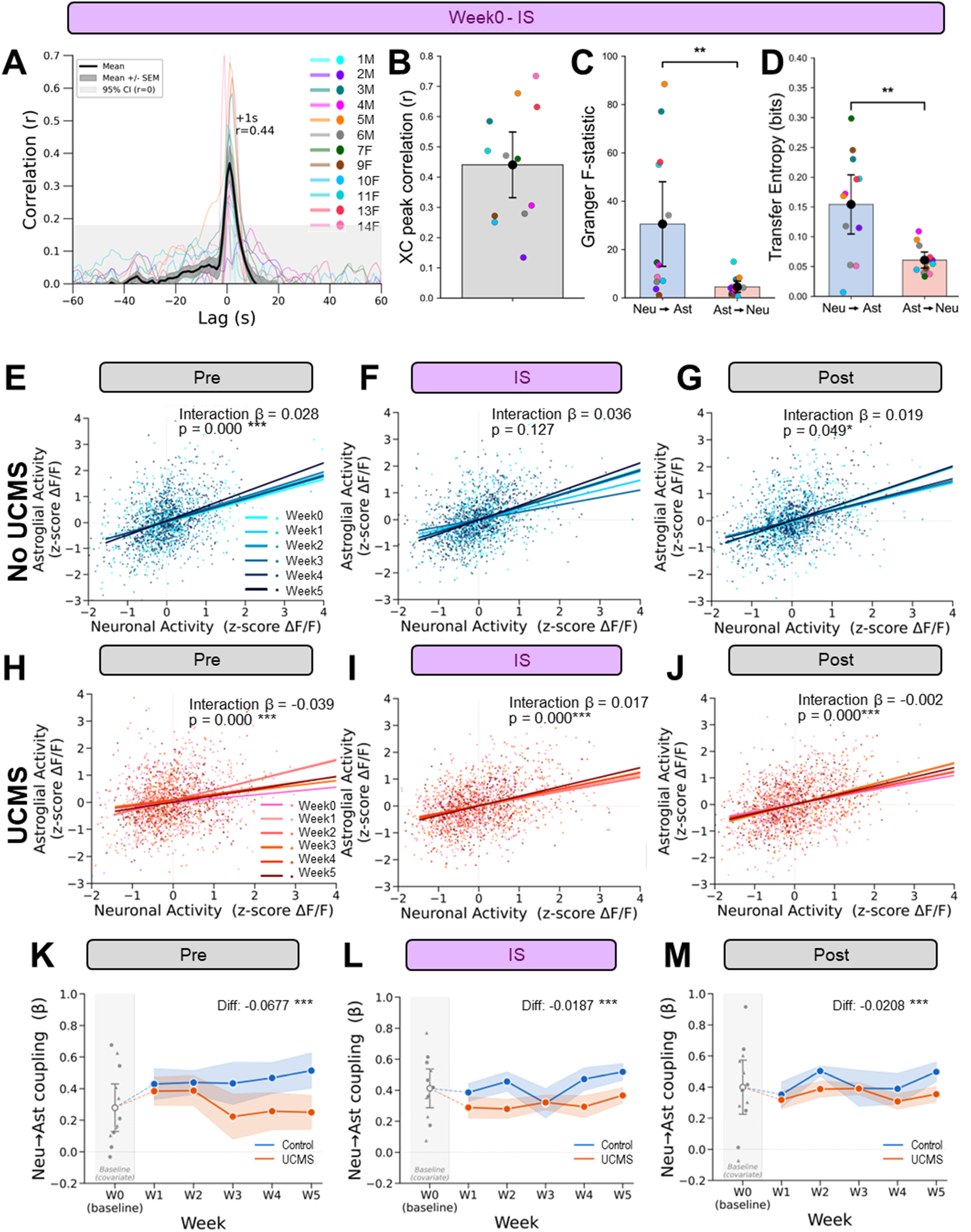
Neuron drive astroglial activity and this process is altered by recurrent IS and UCMS. **(A)** Cross-correlation (XC) between neuronal and astroglial z-scored ΔF/F signals during IS at Week 0. Colored lines show individual subjects, black line indicate group mean and grey band indicate SEM. Grey shaded region indicates the 95% confidence interval for r = 0. Peak lag and peak r of the group mean are annotated. **(B)** Per-animal XC peak correlation values at Week 0 during IS. Histogram represents mean ± s.e.m; dots, individual subjects color-coded as in A. **(C)** Granger causality F-statistic for Neu→Ast and Ast→Neu directions. **(D)** Transfer entropy for Neu→Ast and Ast→Neu directions. **p<0.01. **(E–G)** Scatter plots of astroglial and neuronal z-score ΔF/F in no-UCMS animals across Weeks 0–5 during the Pre **(E)**, IS **(F)**, and Post **(G)** conditions. Lines show linear mixed-model (LMM) fits per week. **(H–J)** Scatter plots of astroglial and neuronal z-score ΔF/F in UCMS animals across Weeks 0–5 during Pre **(H)**, IS **(I)**, and Post **(J)** conditions. **(K–M)** Weekly Neu→Ast coupling strength (slope β from per-week LMMs of Ast ∼ Neu) across Weeks 1–5 for no-UCMS and UCMS animals during the Pre **(K)**, IS **(L)**, and Post **(M)** conditions. Week 0 is shown and was used as a baseline covariate. Diff indicates β differences between no-UCMS − UCMS coupling slopes. *p<0.05, ***p<0.001

### Repeated IS strengthens while UCMS impairs neuron–astroglia directional Ca^2+^ signal coupling across weeks

Having established that neuronal Ca^2+^ signals drive astroglial Ca^2+^ activity, we next examined how this neuronal control evolves across repeated weekly IS challenges, first within no-UCMS and UCMS animals separately across pre-IS, IS, and post-IS conditions (Figure 5E–J), and then between groups (Figure 5K–M). For each condition and group, we fit a linear mixed-effects (LMM) model regressing astroglial z-scored ΔF/F onto concurrent neuronal z-scored ΔF/F, with Week0 as a covariate to obtain resulting slopes Neu-drive across week1-5. In this model Week × Neu-drive interaction captured weekly drift in the directional coupling slope; animal identity was modelled as a random effect to account for repeated within-animal measurements.

In the no-UCMS control group, the within-timepoint relationship between neuronal and astroglial Ca^2+^ activity remained robust across all weeks and all conditions (Figure 5E–G). The Week × Neu-drive interaction was positive and statistically significant during the pre-IS (β=0.028, *p*<0.001) and post-IS (β=0.019, *p*=0.049) conditions, and was not significant during the IS conditions (β=0.036, *p*=0.127), suggesting that neuronal control on astroglial Ca^2+^ activity within this functional coupling remained stable or modestly strengthened with repeated IS exposure in no-UCMS control group.

In UCMS animals, by contrast, the weekly trajectory of coupling diverged across conditions (Figure 5H–J). Pre-IS, the Week × Neu-drive interaction was negative and statistically significant (β=−0.039, *p*<0.001), indicating a progressive weakening or impairment of baseline neuronal drive of astroglial Ca^2+^ activity with cumulative UCMS exposure (Figure 5H). During IS, the interaction was positive (β=0.017, *p*<0.001), suggesting that acute IS challenge transiently increased coupling in UCMS mice each week (Figure 5I). Post-IS interaction was effectively zero (β=−0.002, *p*<0.001), consistent with impaired neuronal control observed in pre-IS condition (Figure 5J).

It is important to note that the LMM β values are small in absolute magnitude but can reach high statistical significance, due to the number of observations/mouse/condition/week (1,800), which contribute to each slope estimates and yield standard errors that are dominated by between-mouse consistency rather than absolute effect size. Significant β coefficients should therefore be interpreted as evidence of reproducible week-over-week trends within each group rather than between weeks quantitatively biologically large effects. Nevertheless, the direction of the relationship between neuronal-astroglial coupling carries biological meaning: in UCMS animals, baseline neuronal control of astroglial activity progressively weakens across weeks (negative pre-IS β), while the IS challenge transiently reverses this trajectory (positive IS β) in both no-UCMS and UCMS animals.

To directly compare the two groups and test whether UCMS affects the longitudinal trajectory of neuron-astroglia functional coupling in a phase-specific manner, we fit a four-way LMM with Group, Condition (pre-IS, IS, post-IS), Week, and their interactions, including baseline coupling at week0 as a covariate to adjust for individual differences before UCMS exposure. Per-mouse random intercepts and Neu-drive slopes were included to allow individual differences in baseline activity and coupling magnitude (Figure 5K–M). In this model, pre-IS served as the reference condition; the Neu-drive x Group x Week coefficient therefore directly tests the group differences in the weekly directional coupling slope at pre-IS, which revealed the largest impairment in neuron-astroglia functional coupling among the three conditions in UCMS animals (Δβ=−0.068, *p*<0.001; Figure 5K). The four-way Neu-drive × Group × Condition × Week interactions then test whether the group divergence at IS and post-IS deviates from this pre-IS reference. Both four-way terms were highly significant (Neu-drive × Group × IS × Week: χ²=58.09, *p*<0.0001; Neu-drive × Group × post-IS × Week: χ²=55.29, *p*<0.0001; joint LRT: χ²=73.24, *p*<0.0001), demonstrating that the weekly trajectory of directional coupling differs between no-UCMS and UCMS animals in a condition-dependent manner. Decomposed per-condition slopes confirmed that UCMS animals showed a flatter or negative weekly trajectory compared to no-UCMS controls, with significant group differences at IS (Δβ=−0.019, *p*<0.001; Figure 5L) and post-IS (Δβ=−0.021, *p*<0.001; Figure 5M). Together our data suggest that UCMS progressively impairs neuronal control of astroglial Ca²⁺ signaling across all three conditions, with the strongest ∼3.5-fold disruption at baseline (pre-IS) and significant but attenuated disruption persisting during and after the acute stress challenge.

Collectively our results demonstrate that whereas recurrent intermittent acute stress exposure enhance neuronal drive on astroglial activity in no-UCMS controls, prior UCMS exposure impairs this process rendering astroglia essentially silent to neuronal control and functionally blunted to acute-stress-evoked Ca^2+^ response.

## Discussion

This study demonstrates for the first time that cortical neuronal and astroglial responses and their functional coupling differs with the duration, recurrence, and chronicity of stress exposure, addressing a fundamental question in neurobiology: how does neural cells interaction evolves during the transition from adaptive to maladaptive stress response? Using dual-color *in vivo* fiber photometry, we mapped PFC Ca²⁺ dynamics across a graded series of stressors. Brief tail pinch and IS increased Ca²⁺ activity in both cell populations, with markedly enhanced coordination during the stimulus that persisted post-stress. When IS was repeated weekly in controls, per-cell-type Ca^2+^ changes remained stable while functional coupling between neurons and astroglia progressively strengthened across pre-IS, IS, and post-IS conditions, suggesting adaptive response to stress. In UCMS animals, this recurrent IS response trajectory diverged sharply by week 5: neuronal IS reactivity was approximately doubled, astroglial activity was blunted, and neuron-astroglia coupling collapsed with a ∼3.5-fold disruption at baseline (pre-IS) and significant, though attenuated, disruption during and after IS. This coupling breakdown was associated with loss of neuronal drive on astroglial activity and emerged in parallel with the onset of UCMS-induced anxiety- and anhedonia-like deficits.

### Acute stressors elicit coordinated neuronal and astroglial responses in the PFC

Our finding that brief tail-pinch stress elicits coordinated Ca²⁺ transients in PFC neurons and astroglia is consistent with prior single-cell-type recording studies of nociception or stress in either population individually [56–58]. Here using both single-color and dual-color recordings, we confirmed these results in the PFC and extended them by demonstrating that the two cell populations are engaged simultaneously and become more tightly coupled during the stimulus. Similar tail-pinch-evoked neuronal Ca²⁺ responses have been reported in other brain regions [57, 59, 60] and in neurons projecting to the PFC [61]. Our astroglial findings also converge with work by Pittolo *et al.* [62], showing that tail-pinch-evoked astroglial Ca²⁺ elevations in the PFC are mediated by dopamine and noradrenaline via α1-adrenergic receptors. Mechanistically, we found that stimulus-locked Ca²⁺ signals emerged in both cell types, with a brief lag (∼0.3 s) in the astroglial Ca^2+^ wave, suggesting localized astroglial recruitment driven by upstream neuronal signaling. To our knowledge, only one prior study by Liu *et al*. [63] has reported comparable delay between neuronal and astroglial response, showing habenular Ca^2+^ changes following foot shock, broadly consistent with the delay we observed in the PFC. Functionally, our findings of astroglial recruitment within seconds of an acute stressor adds to growing evidence that PFC astroglia is engaged when neuronal activity elevates during behavior such as anxiety [46]. Several mechanisms could mediate this rapid recruitment, e.g. activation of astroglial membrane receptors and transporters that drive intracellular Ca²⁺ signaling. These mechanisms may include α1-adrenergic receptors as aforementioned [49], and/or local increases in glutamatergic transmission at the tripartite synapse [38, 64, 65] likely converging on IP₃R2-dependent Ca²⁺ release from internal stores, the somatic signal preferentially captured by fiber photometry [66].

Immobilization or restraint stress is a widely used laboratory paradigm to investigate stress-related physiological, cellular and behavioral changes [67]. Restricting animal movement on a flat platform enables continuous Ca²⁺ recording in cannulated animals while inducing plasma corticosterone elevations comparable to those produced by tube- or mesh-based immobilization [67]. Prior fiber photometry studies have shown that restraint stress drives Ca²⁺ activity in neuronal populations across several stress-sensitive circuits, including the insula–BNST [68], the hypothalamus-habenula [60] and lateral septum and extended circuits [45, 69]. Here, we expand these findings to PFC neurons, converging with prior c-Fos immunostaining and brain-wide cell activity mapping studies that consistently reported PFC neuronal activation by acute stress [27, 32, 70]. Importantly, here we demonstrate that astroglia are not passive bystanders but are recruited by acute stress paralleling recent c-Fos brain-mapping evidence of stress-evoked astroglial activity [27]. Beyond confirming individual cell-type activation, our dual-color approach reveals a cellular dimension previously inaccessible to single-population methods i.e. intercellular relationship during sustained acute stress. Specifically, neuronal and astroglial Ca²⁺ event frequencies were positively correlated during IS and remained elevated post-stress, with similar coupling observed during and after tail pinch. It is important to mention that we found differences between pre-pinch and pre-IS coupling strength. Indeed, pre-pinch within-animal Ca^2+^ signal coupling was detectable over a 5 sec windows on continuous Ca^2+^ fluctuations (50 data point per sec). However, no comparable pre-IS coupling was observed at discrete event-frequency level (1 datapoint per min). This apparent difference is not a discrepancy, but rather reflects the analytical resolution, and is consistent with the expectation that spontaneous Ca^2+^ events during the 30 min window occur asynchronously between cell types in the absence of an external stimulus.

Another informative aspect of our data concerns the post-stress condition. While neuronal Ca²⁺ responses to IS persisted beyond the stressor exposure, astroglial Ca²⁺ elevation subsides, yet functional coupling between the two cell types remained strong. Sustained post-stress neuronal activity has previously been described in the BNST and proposed as a substrate for stress resilience since blocking this post-stress signaling conferred vulnerability and precipitated the emergence of depressive-like behavior [44, 71]. Our results add a new dimension to this picture; and suggests that in the PFC, what persists post-stress is not only the elevated neuronal activity but a strengthened functional partnership between neurons and astroglia.

### Repeated intermittent homotypic stress enhance neuronal and astroglial activity and functional coupling

A key strength of our experimental design was that the control animals received repeated weekly IS challenges in parallel with the UCMS group, allowing us to characterize the adaptive response to recurrent homotypic stress in animals with no prior UCMS exposure. We found that while the first IS challenge drastically increased Ca^2+^ transients in neurons, IS response sightly subsides with following exposures (2-fold vs 50-60%). Stable and relatively more modest increases in astroglial Ca^2+^ event rates were observed across challenges. Overall neuronal and astroglial Ca^2+^ responses were largely similar across recurrent homotypic stress. We might have expected neuronal or astroglial habituation or adaptation [4, 64] e.g. waning of individual cell population Ca^2+^ signals, which we did not observe. Yet when we examined intercellular functional coupling, a clear adaptive signature emerged. We observe this with *rmcorr* analysis where the weekly correlation coefficients (r) indexing neuron-astroglia coupling progressively strengthened across weeks, pre-, during- and post-IS. We also found that the neuronal drive on astroglial activity increased with time pre- and post-IS (positive β). Both findings suggest that functional coupling within the neuron-astroglia network, rather than the activity of either cell type alone, may serve as a more sensitive index of stress adaptation with enhanced cell-cell communication across stress phases, and tightened neuronal control of astroglial activity during the pre- and post-stress. These two complimentary processes may play a key role in adaptative response to stress.

### UCMS impairs neuron-astroglial functional coupling and neuronal drive on astroglia

Chronic unpredictable stress rodent models are numerous and vary depending on the severity, nature, duration of different stressors used as well as the length of the chronic stress exposure [72, 73]. Here we adapted our classical UCMS protocol to allow weekly recordings during IS and showed that behaviorally, the modified protocol reproduced the hallmark features of chronic stress [51, 52, 55] emerging by weeks 4–5, confirming that fiberoptic implantation and repeated recording did not prevent the detection of the development of UCMS-induced anxiety- and anhedonia-like deficits. We assessed how chronic UCMS altered the longitudinally neuronal and astroglial Ca^2+^ activity at baseline and in response to IS. Chronic stress produced a cell-type-specific divergence in the response to the IS challenge: neuronal Ca²⁺ reactivity was sensitized, with the IS-evoked increase in event frequency roughly twice that of no-UCMS controls, whereas the astroglial IS response was blunted. The sensitized neuronal response is consistent with the broader view that prior chronic stress renders cells of the PFC hyperexcitable to subsequent stressor. Indeed, we previously reported that chronic restraint stress and UCMS enhanced acute-stress-evoked neuronal activation in the PFC, indexed by c-Fos, with the magnitude of this exacerbation positively correlating with the severity of chronic stress-induced behavioral deficits [74]. The present study extends this finding to neuronal Ca^2+^ activity. The progressive nature of the changes is in line with work showing that chronic stress incrementally potentiates stress-relevant circuitry [60] and may be a key process involved in maladaptive response to stress.

In contrast to the sensitized neuronal response, prior UCMS exposure progressively blunted the astroglial Ca^2+^ response to IS. Indeed, no-UCMS controls displayed a reliable 30-59% IS-evoked rise in astroglial Ca^2+^ activity, whereas by UCMS end, the IS-evoked response was essentially absent. This blunting was not attributable to UCMS-induced astroglial loss, as the number of recorded cells did not differ between groups and the same pattern remained when responses were normalized to each animal’s own baseline. The blunted astroglial Ca²⁺ response is consistent with well-documented astroglial dysfunction under chronic stress. Studies, including our own previous work, have reported reduced astroglial marker expression [16], impaired metabolism [75], and morphological atrophy in PFC astroglia following chronic stress [12, 14, 76]. Our recordings demonstrates that these molecular and structural changes are associated with impairment of astroglial cell activity. These astroglial molecular and cellular changes closely parallel those observed in post-mortem findings in depression [13, 14, 17, 20], raising the possibility that the astroglial abnormalities observed in depression reflect a stress-induced impairment of cortical astroglia function.

Importantly, the simultaneous sensitization of neurons and hypo-responsivity of astroglia correspondingly weakened their intercellular relationship. By examining how neurons drive astroglial activity longitudinally under chronic stress, we uncovered a previously unaddressed form of neuron-astroglia circuit pathology where UCMS progressively uncoupled the astroglial from their local neuronal partners. This trajectory is the reverse image of the adaptive response observed under repeated IS in no-UCMS controls with weakened Ca^2+^ coupling in pre- and post-IS conditions. The impaired intercellular coordination likely involves multiple mechanisms, including impaired glutamate clearance via reduced glial glutamatergic transporter-1 (GLT1 or human homologue EAAT2) expression [77], disrupted potassium buffering [78] and altered metabolic coupling through the astrocyte-neuron lactate shuttle [79], all of which have been implicated in chronic stress vulnerability. We therefore interpret the cell-type-specific divergence not as two independent changes but as a breakdown of the coordinated neuron–astroglia operation that normally constrains cortical excitability under stress. Indeed, an inability of PFC astroglia to modulate their own activity in response to neuronal drive would result in hyper-sensitized neurons, likely driving the transition from adaptive to maladaptive state in the cortex and its downstream targets. Critically, the timing of this coupling collapse may precede or aligns with the emergence of behavioral deficits where loss of coupling became detectable around weeks 2–3 and the loss of neuronal drive around weeks 4-5. This is consistent with the possibility that progressive circuit-level reorganization between neurons/synapses and astrocytes contribute to, rather than merely accompanies, the transition to a depressive-like state [16, 60].

Several limitations of this study should be acknowledged. Although fiber photometry provides a powerful *in vivo* readout of cell-type-specific activity and intercellular coupling, it carries inherent constraints. Notably, comparing baseline Ca²⁺ activity across animals is challenged by variability in viral infection rates and small fluctuations in signal stability across longitudinal recordings. We addressed this through careful experimental design, within-animal normalization, and the use of within-subject statistical approaches (*rmcorr* and LMM with per-animal random effects). Although we found no sex differences in tail-pinch- or IS-evoked responses, the UCMS cohort was not powered to formally test sex as a factor in the analyses of recurrent acute stress or chronic stress responses; future studies should examine sex-specific effects and the potential contribution of neuron-astroglia coupling to sex-related vulnerability to stress. Our approach also did not resolve coupling between astroglia and specific neuronal subtypes (e.g., pyramidal vs interneuron populations) or across brain regions, nor did we test whether antidepressants restore functional coupling lost under UCMS, cell-type-resolved investigations combined with pharmacological or genetic rescue represent natural next steps for testing causal engagement and therapeutic tractability.

Together, these findings establish neuron-astroglia functional coupling as a dynamic cellular substrate distinguishing adaptive from maladaptive stress responses. Acute, sustained, and recurrent stressors enhanced coupling strength and neuronal drive, identifying functional coupling as a more sensitive index of adaptation than individual cell-type activity. Under chronic stress, this coordination collapsed as PFC neurons became hypersensitized, astroglia became hyporesponsive to neuronal drive, and the two populations functionally uncoupled prior or when with depressive-like behavior deficits emerge. Our findings position the loss of neuron-astroglia functional coupling as a potential early substrate for stress-induced cortical dysfunction and a mechanistically tractable therapeutic target for stress-related psychiatric disease.

## Materials and methods

### Animals

Mice (8-week-old; C57/Bl-6, Jackson Laboratories, lot # 000664, 50% female) were housed under standard conditions at a constant temperature of 20–23°C on a 12-h light/dark cycle. After surgeries, mice were single housed for rest of the procedures, with ad libitum access to food and water except when required for UCMS stressors or behavioral testing. All procedures were approved by the Centre for Addiction and Mental Health Centre for Addiction and Mental Health (CAMH certificate #0038; AUP #2021-857) and followed the guidelines of the Canadian Council on Animal Care (CCAC).

### Viral infusions and cannula implantation

As illustrated in experimental design (Figure 1A), mice were anaesthetized with isoflurane and unilaterally infused with adeno-associated viruses (AAVs) expressing genetically encoded Ca^2+^ indicators in the PFC (PFC; AP +2.0, ML ±0.3, DV −3.0 relative to bregma) using stereotaxic surgery. For single-channel recordings, astroglial Ca²⁺ activity was monitored using pZac2.1 gfaABC1D-cyto-GCaMP6f-AAV5 (Addgene #52925, titer ≥ 7×10^12^ vg/mL, n = 8) or neuronal Ca²⁺ activity using pAAV5-SYN1-GCaMP6f-WPRE-SV40 (Addgene #100837, titer ≥ 7×10^12^ vg/mL, n = 8). For simultaneous dual-color recording, mice were co-infused with pZac2.1 gfaABC1D-cyto-GCaMP6f-AAV5 and pAAV5.Syn.NES.jRCaMP1a.WPRE.SV40 (Addgene #100848, titer ≥ 1×10^13^ vg/mL at a 1:1 ratio (n = 16). Viral suspensions were delivered at 0.1 µL/min (single-channel: 0.5 µL total volume; dual-color: 1.0 µL total), and a monofiberoptic cannula (400 µm core, 0.48 NA; Doric Lenses, Quebec, Canada) was implanted 200 µm above the injection site. Half of the animals were infused/implanted on the right PFC and the other half in the left. Animals received post-operative analgesia and were monitored daily for signs of distress for 3 days. Cohort sizes for the dual-color experiments were reduced following surgery: two animals lacked analyzable signals on both channels and two lost their cannulas during the immobilization stress (IS) protocol, yielding final samples of n = 14 for the tail-pinch experiment and n = 12 for the remaining dual-color recordings.

### Acute stress challenges

After ≥ 4 weeks of recovery and stable transgene expression, mice were habituated to the recording room for 1 hr and to the patch cord for an additional 15 min before each session.

#### Tail-pinch challenge

Mice were gently held in the palm of the hand for 2 min, after which a 400-g pressure was applied 1 cm above the tip of the tail for 3 s using a Rodent Pincher Algometer (Bioseb, France). Recording continued for a further 2 min following stimulus offset (Figure 1B) [58, 80, 81].

#### Immobilization stress (IS) challenge

Each session comprised three continuous 30-min phases: a baseline period in the home cage (pre-IS), an immobilization period in which mice were placed in an empty cage without bedding and gently secured with sensitive-skin tape to restrict limb and body movement, and a recovery period (post-IS) during which the tape was carefully removed and mice were returned to their home cage (Figure 1C) [82].

### Unpredictable chronic mild stress (UCMS) and behavioral testing

Mice were exposed to UCMS for 5 weeks, receiving 2–3 randomized stressors per day in an unpredictable schedule. Stressors included predator odor, light-cycle disruption (4 h of darkness during the day or constant overnight illumination), cage tilting (45°), reduced cage space, and removal of bedding or nestlet. The protocol was adapted from previous work [20, 51, 52] but omitted stressors incompatible with chronic fiber-optic implantation, including those that could promote wound infection (e.g., forced bath, wet bedding) or directly confound the immobilization challenge (e.g., restraint). Control (no-UCMS) animals were maintained under standard housing throughout. To verify that the adapted protocol reproduced previously characterized behavioural deficits [12, 35] both groups underwent weekly behavioural testing using the PhenoTyper test [12, 54, 76] and the sucrose consumption test [12, 76], and coat-state degradation [12, 54, 76]. Behavioural testing procedures were identical to those described previously [12, 76] (full procedures in Supplementary Methods). Briefly, anxiety-like behaviour was assessed in home cage-like apparatus (Noldus® phenotyper) in which a 1 hr (11:00 pm – 12:00 am) light challenge was applied over the food zone. Shelter-zone time over the subsequent 5 hrs (12:00 – 5:00 a.m.) was normalized to the no-UCMS group mean and expressed as residual avoidance behavioral index [16, 54]. The sucrose consumption test which measures anhedonia-like behaviour which consists in habituation to 2 % sucrose solution before UCMS onset and weekly a 1 hr sucrose consumption test (2–3% sucrose) preceded by 16 h of fluid deprivation, with water consumption measured weekly under matched conditions. Coat-state deterioration was scored weekly across 7 body parts [12, 54, 76].

### In vivo fiber photometry recordings and analysis

#### Preprocessing

Ca^2+^ signals were acquired using Doric fiber photometry systems (Basic FPS – GCaMP + Red Fluo; Doric Lenses, Quebec, Canada). For single-channel recordings, excitation was delivered alternately at 405 nm (isosbestic control) and 470 nm (GCaMP6f signal). For simultaneous dual-color recordings, 470 nm and 580 nm LEDs were used to excite GCaMP6f and jRCaMP1a fluorophores, respectively and normalized to isosbestic channel, enabling concurrent measurement of astroglial and neuronal Ca²⁺ activity. Fluorescence emission signals were collected at 12 kHz, and laser intensity at the fiber tip was kept low to minimize photobleaching. Signals were down-sampled to 100 Hz, and both the activity and isosbestic channels were filtered to remove brief electrical spikes and high-frequency noise. Slow photobleaching drift was estimated and subtracted from each channel using airPLS, and movement artifacts were corrected by scaling the isosbestic channel to the signal channel and subtracting it, yielding ΔF/F. For tail-pinch-evoked response analysis, ΔF/F was z-scored to a 5 s pre-pinch baseline (the 5 s immediately preceding stimulus onset) so that all values within the analysis window (5 s baseline, 3 s pinch, 17 s post-pinch) reflected changes relative to the pre-pinch condition. For the IS and UCMS experiments, ΔF/F was z-scored within each weekly recording to enable comparison across animals and sessions. The z-scored traces were then used for event detection (yielding per-minute event-frequency time series) and for all downstream directionality and mixed-effects analyses.

#### Quantification of Ca2+ activity

For both single-channel and dual-color recordings, tail-pinch-evoked responses were quantified using z-scored ΔF/F (referenced to the 5-s pre-pinch baseline) and reported as mean ± SEM across animals. The area under the curve (AUC) was calculated for each time window (pre-, pinch, post-) using the trapezoidal method. For dual-color recordings, the temporal lag between neuronal and astroglial Ca²⁺ signals were estimated for each animal using the maximum cross-correlation method [43], providing the first quantitative measure of directional coupling between the two cell types. For IS-evoked responses, analyses were performed primarily on Ca²⁺ event frequency and peak amplitude. Ca²⁺ events exceeding 3× the median absolute deviation (MAD) of the z-scored ΔF/F trace for neurons [83] and 2.5× MAD for astroglia [84] were quantified, with peak amplitude calculated relative to the local baseline. Event frequency was expressed as events per minute, and peak amplitude as ΔZ-scores. See Supplementary Methods for details.

### Analysis of neuron–astroglia directionality and coupling

Several complementary methods were used to identify changes in neuron-astroglia functional coupling, all based on correlations between neuronal and astroglial Ca²⁺ signals. For the tail-pinch experiment, rapid changes in temporal coupling were assessed using cross-correlation analysis within each window (pre-, pinch, and post-pinch), with Pearson correlation assessing individual-animal signal relationship and repeated-measures correlation (*rmcorr*) estimating the common within-subject slope across the cohort [50]. These analyses were applied to the z-scored ΔF/F traces (50 datapoints per second) for the tail-pinch challenge or per-minute event frequencies for the IS challenge, to evaluate the coupling strength individually for each of the three windows/conditions (pre-, during, and post-acute stress). The same per-window *rmcorr* was performed weekly in either no-UCMS or UCMS groups to track progressive IS coupling changes in each group.

Because *rmcorr* does not allow direct comparisons between groups or time points and does not address the direction of information flow, we then performed three parallel analyses (cross-correlation lag [85, 86], Granger causality [87], and Transfer Entropy [88, 89]) to determine the temporal directionality of coupling between PFC neuronal and astroglial Ca²⁺ signals, followed by a linear mixed-effects model (LMM) [90] to infer weekly coupling trajectories and quantify between group differences.

Cross-correlation lag, Granger causality, and Transfer Entropy are complementary methods that capture distinct aspects of directionality: temporal lag between signals (cross-correlation), linear predictive influence under autoregressive modelling (Granger), and both linear and nonlinear predictive dependencies (Transfer Entropy). These analyses were performed on neuronal and astroglial z-scored ΔF/F traces from the week-0 IS recording, resampled to 1 Hz. using. Briefly, cross-correlation was obtained across a ±60-s lag ranges with the peak position indicating the time offset of maximal alignment between the two cell-type signals. Granger causality was tested using linear ordinary least squares autoregressive models comparing F-statistics between directions (Neu→Ast vs. Ast→Neu) to test whether 10s-one signal’s past improved prediction of the other beyond its own history. Transfer Entropy was estimated using a histogram-based discrete-state estimator with 6s-history length and 8 quantile bins, with directionality summarized as the asymmetry index (TE_Neu→Ast − TE_Ast→Neu) / (TE_Neu→Ast + TE_Ast→Neu). For each method, the two directions were compared within-animal across the cohort using paired Wilcoxon signed-rank tests.

All directionality analyses were implemented in Python (NumPy and SciPy): Granger causality used numpy.linalg.lstsq for ordinary least squares estimation of the autoregressive models and scipy.stats.f for *F*-test *p*-values, while Transfer Entropy was computed from histogram-based Shannon entropy estimates using numpy.percentile for quantile binning and numpy.unique for joint-distribution counting. Cross-correlation was computed using numpy.corrcoef evaluated at each integer lag.

We then used LMM as statistical framework enabling the comparison of timing and magnitude of signals across conditions while accounting for between-animal differences, an approach previously used for fiberphotometry-based Ca^2+^ recording [91]. Here, a LMM was fit to characterize how the within-animal relationship between neuronal and astroglial Ca²⁺ activity evolves across conditions (pre-IS, IS and post-IS), timepoints (weeks 0 to 5) and across groups (no-UCMS vs UCMS). The LMM was carried out as a custom maximum-likelihood estimator using scipy.optimize.minimize with the Nelder-Mead algorithm; likelihood ratio tests were computed using scipy.stats.chi2. The model regressed astroglial z-scored ΔF/F on concurrent neuronal z-scored ΔF/F (the resulting slope hereafter referred to as Neu-drive) across weeks 1–5 (1,800 observations/mouse/condition/week), with fixed effects for group, condition (with pre-IS as reference), week, and all relevant interactions; animal identity was modelled as a random effect. Within-condition week0 coupling values were included as a covariate to adjust for pre-UCMS individual differences. Two nested specifications were fit with increasing levels of interaction complexity: Model A isolated all main effects and three-way interactions including Neu-drive × Group × Week, while Model B additionally included the two four-way interactions Neu-drive × Group × Condition (IS) × Week and Neu-drive × Group × Condition (Post) × Week to test whether the group differences in the weekly coupling slope varied by condition. Fixed effects were tested with Wald z-statistics, and nested models were compared using likelihood ratio tests.

### Immunohistochemistry

Within 24 hr of the final fiber photometry session, mice were anaesthetized with avertin (125 mg/kg, i.p.) and transcardially perfused with 4% paraformaldehyde (PFA, pH 7.4). Brains were post-fixed overnight at 4 °C in 4% PFA, cryoprotected in 30% sucrose for 48 h, and frozen on dry ice. Coronal PFC sections (20 µm) were cut on a cryostat (Leica CM1950), mounted on charged slides, and stored at −80 °C until use. Sections were immunostained to verify cannula placement and AAV specificity using cell-type-specific markers (NeuN for neurons, GFAP for astroglia) and sensor-specific markers (anti-GFP and anti-RFP antibodies). Images were acquired on Olympus IX83 inverted microscope from 3–4 coronal sections per animal spanning the viral infusion site (AP +2.0 to +2.5 mm relative to bregma). Viral infection rates were quantified in QuPath (version 0.5.0), with the percentages of GFP⁺/GFAP⁺ astroglia and RFP⁺/NeuN⁺ neurons calculated relative to the total GFAP⁺ and NeuN⁺ cell populations, respectively. The full antibody panel and staining protocol are provided in the Supplementary Methods.

### Statistical analysis

For tail-pinch- and IS-evoked Ca²⁺ responses, repeated-measures ANOVA was performed with time and sex as factors, followed by Tukey’s *post hoc* test to compare z-scored ΔF/F, AUC, maximum cross-correlation, event frequency, and event amplitude across the pre-, during-, and post-stimulus conditions. The similar analysis was performed separately within each weekly recording and within each group to assess the per-week effect of acute IS challenge. Although both male and female animals were included in the UCMS cohort, the sample size was not sufficient to support sex as a factor in the statistical analyses; data were therefore collapsed across sex. Progressive changes across weeks in coat-state, RA, and sucrose consumption were tested using repeated-measures ANOVA with week and group as factors, followed by Tukey’s *post hoc* tests. To compare neuronal and astroglial Ca²⁺ changes between groups, two-way ANOVA was applied to Ca²⁺ event frequencies expressed as percent change from each group own baseline, with group and IS condition as factors. All ANOVA-based analyses were performed in GraphPad Prism (version 10); significance was set at p < 0.05, and data are presented as mean ± SEM. Statistical procedures specific to correlation, directionality, and LMM analyses as well as software used are described in their respective sections.

## Supporting information

Supplementary Method and Results

## Acknowledgements

We thank the CAMH animal facility staff: Kristen Fournier, Katrina Deverell, and Taylor Pacheco for their assistance with animal care. We also thank Rosemary Bagot and Jessie Muir for their initial help in setting up our lab fiberphotometry platform.

## Generative AI statement

Anthropic’s Claude (4.6 and 4.7) and Google’s Gemini 3.5 were used for Python code assistance during data analysis and for readability and language refinement of the final manuscript. After the application of AI, the authors reviewed and edited the manuscript to ensure its content accuracy.

## Conflict of Interest

The authors declare no competing interests.

## Funding

This work was supported by the CAMH Discovery Fund (PI: M.B.), the Canadian Institutes of Health Research (CIHR; PGT165852, PI: M.B.), and the Campbell Family Mental Health Research Institute. Y.B. was supported by a CIHR Postdoctoral Fellowship (202110MFE-472592-FPP-CEAH-93191), the CAMH Discovery Fund Fellowship (2021-0859), the CAMH womenmind Fellowship (2021-0863) and UofT Labatt Family Research Fellowship in Depression Biology (2021-1001013).

## Author contributions

Y.B. performed experiments, fiber photometry recordings and data analysis. S.A.C. and J.K.K. assisted with animal handling and the stress paradigms. K.C. and Y.S.N. contributed to the correlation analyses. M.A.M contributed to histology and infection rate quantification. Y.B. and M.B. wrote the manuscript. E.S. provided scientific feedback and edits on the manuscript. M.B. conceived and designed the study, supervised the work, and acquired funding. All authors reviewed and approved the final manuscript.

## Code availability

The custom code used for signal processing and associated statistical analysis is publicly available at https://github.com/bansal-yashika/Fiber-photometry-analysis-pipelines.

## Notes

### Competing Interest Statement

The authors have declared no competing interest.

### Summary of Updates

This version of the manuscript has revised figures, manuscript edits, and a new complementary analysis that needs to be discussed in the discussion section.

