## Supplementary Method and Results for "*In vivo* cortical neuron-astroglial functional coupling strengthens with acute stress but is impaired by chronic stress in mice"

**Word count: 2115**

**Number of Supplementary Figures:** 6

**Number of Supplementary Tables:** 3

**Number of Supplementary References:** 5

**SUPPLEMENTARY METHODS**

***Behavioral Assessments:***

*Sucrose consumption test.* This test was used as a measure of anhedonia-like behavior. During the week 0, animals were habituated to a 2% sucrose solution for 48 hrs. Sucrose intake was then measured over a 1 hr session (10:00 – 11:00 am) following 16 hrs of fluid deprivation, after which water access was restored for 48 hrs. The same procedure was then applied to measure water intake (16 hrs fluid deprivation followed by 1 hr water access). In subsequent weeks, mice were re-habituated to sucrose for 24 hrs and retested under the same conditions. Because sucrose intake remained low in both groups through week 3, likely reflecting reduced consumption associated with surgery and repeated weekly recordings, the sucrose concentration was raised to 3% at weeks 4 and 5 to enhance dynamic range and improve sensitivity without changing the test structure. All sucrose and water tests were performed at least 18 hrs after the last UCMS stressor which was the same every week.

*PhenoTyper Test (PT):* We used the PhenoTyper test as a measure of conflict-induced anxiety-like behavior, an assay that produces reliable and reproducible readouts across multiple weeks of testing in mice subjected to chronic stress. The test utilizes the PhenoTyper (Noldus, Leesburg, VA, USA) apparatus, which consists of a home cage like arena (30x30cm) with a shelter, food, and water zone. Mice were placed in the apparatus once weekly during the dark phase (7:00 p.m. – 7:00 a.m.), at least 1 hr after the last UCMS stressor of the day and on the evening following the water-intake test, with timing maintained consistent across weeks. The conflict occurs when a light is automatically switched on over the food zone at 11pm for 1 hr causing the mice to hide in their shelter. Mice subjected to chronic stress including UCMS were shown to display anxiety-like behavior in this test, by continuing to hide in the shelter and avoid the remaining areas of the arena even after the light turns off. Residual avoidance (RA) was computed for each animal using the equation defined in [1]. Briefly, the equation quantifies the time spent in the shelter zone during the 5 h following light offset, normalized to the no-UCMS group mean. By this definition, control animals show RA values of ~0 ± SEM, while UCMS-exposed animals show positive RA values, with higher values reflecting greater anxiety-like behavior. RA can equivalently be computed from avoidance of the food zone but typically mirrors shelter-zone data therefore to avoid redundancy, only shelter-zone measurements are shown in this manuscript.

***Fiber photometry signal processing and data collection***

Signals were down-sampled from 12 kHz to 100 Hz for analysis. Both the activity signal and the 405-nm isosbestic control channel were filtered to remove brief electrical spikes and high-frequency noise. Photobleaching was corrected by estimating the slow downward drift in each channel using adaptive iteratively reweighted penalized least squares (airPLS) and subtracting it. Movement artifacts were then removed by scaling the control channel to match the signal channel and subtracting it. ΔF/F was calculated from dividing the corrected signal by its estimated baseline and converted to z-scores within each recording so that activity could be compared across animals and sessions. For the tail-pinch challenge, the evoked response was instead measured directly from baseline-normalized ΔF/F rather than from detected Ca^2+^ events: the 5 s immediately preceding the pinch were used as the reference (mean and SD) to z-score the 25-s response window (5 s baseline, 3 s pinch, and 17 s post-pinch), so that values reflect change relative to the pre-pinch state. For IS similar z-scoring was performed before event detection and for the directionality and mixed-effects analyses. Specifically, Ca^2+^ events were identified from z-scored traces using SciPy’s find_peaks. The detection threshold was set adaptively from each recording’s own noise level, measured using the median absolute deviation (MAD). The threshold was set at the median plus k × MAD. Because the two cell types differ in signal shape, distinct multipliers were used: *k* = 3.0 for neurons, which show sharp, large transients, and *k* = 2.5 for astroglia (gfaABC1D-GCaMP6f), which show broader, smaller ones. To avoid peak overestimation, each event was also required to meet a minimum prominence (1.0 z-units for neurons; 0.8 for astroglia) and a minimum width (100 ms for neurons; 250 ms for astroglia). The same neuronal parameter set was applied to both neuronal indicators, Syn1-jRCaMP1a (dual-color recordings) and Syn1-GCaMP6f (single-channel recordings).

For each event, peak amplitude was measured relative to a local baseline defined as the median of the 5 s immediately preceding the event. Event frequency per minute and per-event peak amplitudes were averaged within each phase where bins with no events were kept as zeros rather than excluded. Event frequency and amplitude values were used for pre-IS, IS and post-IS ANOVAs for no-UCMS and UCMS groups, and event frequency was used for within group and within-subject repeated-measures correlations (rmcorr).

Data processing and analysis code is available at <https://github.com/bansal-yashika/Fiber-photometry-analysis-pipelines>

**Directionality and longitudinal modelling of neuron–astroglia functional coupling**

The directional relationship between neuronal and astroglial Ca^2+^ signals was examined on the z-scored ΔF/F traces resampled to 1 Hz at week0 during IS. This recording period was selected because acute IS reliably engages both cell populations driving coordinated transients across neurons and astroglia — producing the dynamic, event-rich signals required to reliably estimate directional coupling. Week 0 pre-IS recordings, by contrast, are dominated by low-amplitude spontaneous fluctuations that yield poor signal-to-noise for directionality estimation.

***Cross-correlation***: Cross-correlation between concurrent z-scored neuronal and astroglial ΔF/F traces was computed as a Pearson correlation evaluated on overlapping samples at integer lags from −60 to +60 s; yielding a correlogram that describes how strongly the two signals align as a function of temporal offset k [2]. By convention, positive lags indicate that neuronal signal precedes astroglial signal (i.e., the neuronal signal at time t predicts the astroglial signal at time t + lag), while negative lags indicate that astroglial signals leads. The lag at peak correlation for each sampling window was used to calculate average lag between signals for each animal, and the correlation amplitude at that lag (r_max) served as a scalar measure of coupling strength. Correlations exceeding the 95% null envelope of ±1.96/√N (where *N* is the number of samples per recording) were considered significantly different from zero.

***Granger causality:*** This method test whether neuronal Ca^2+^ signal predicts future astroglial Ca^2+^ signal beyond what astroglial history alone can account for (and vice versa). Bivariate Granger causality was estimated between the two z-scored ΔF/F signals using linear ordinary least-squares (OLS) autoregressive models with a fixed lag order of 10 samples (= 10 s at 1 Hz). For each directional test, two nested autoregressive models were fit to the target signal. For the Neu → Ast direction, the restricted model regressed the astroglial signal at time *t* on its own past 10 samples (Ast[*t* − 1] through Ast[*t* − 10]); the full model additionally included the neuronal signal's past 10 samples (Neu[*t* − 1] through Neu[*t* − 10]) as predictors. The reverse Ast → Neu direction was tested symmetrically, with the neuronal signal as the outcome. F-statistics was calculated from residual sum of squares from the restricted to the full model; a significant *F* indicates that the predictor signal's past carries predictive information about the outcome signal's future beyond what the outcome's own past provides. *F*-statistics from the two directions were compared across the cohort using a paired Wilcoxon signed-rank test.

***Transfer entropy (TE)***: Because Granger causality assumes linear Ca²⁺ dynamics and may underestimate nonlinear coupling, we complemented it with TE, a model-free measure that captures both linear and nonlinear directional dependencies between signals. TE was calculated from the two z-scored ΔF/F traces after discretization into eight equiprobable bins in line with standard practice for histogram-based TE estimation of moderate-length time series [3]. A 6-s history window was used for each signal, spanning the typical decay timescale of Ca²⁺ transients (~3–5 s for neuronal events; ~5–8 s for astroglial events) and approximately matching the autocorrelation decay of both signals. Conceptually, TE quantifies in bits the extent to which knowing the neuronal signal's recent past reduces uncertainty about the astroglial signal's next value beyond what the astroglial signal's own past already provides. In other words, a non-zero value indicates that the neuronal signal carries predictive information about future astroglial activity beyond what astroglial autocorrelation alone explains. The TE of reverse direction (Ast → Neu) was quantified symmetrically. To summarize the net direction of information flow per animal, a asymmetry index (AI) was calculated as the difference between the two directional TE values divided by their sum, resulting in AI ranging from −1 to +1, with positive AI values indicating predominantly neuron-led coupling. The two TE directions were compared within-animal using paired Wilcoxon signed-rank tests. Both TE’s are reported in the Figure 5 and AI in the main text.

All directionality analyses were implemented in Python (NumPy and SciPy): Granger causality used numpy.linalg.lstsq for OLS estimation of the autoregressive models and scipy.stats.f for *F*-test *p*-values, while Transfer Entropy was computed from histogram-based Shannon entropy estimates using numpy.percentile for quantile binning and numpy.unique for joint-distribution counting. Cross-correlation was computed using numpy.corrcoef evaluated at each integer lag.

***Immunohistochemistry: detailed protocol and antibodies***

For immunohistochemistry, animals were perfused under anesthesia (Avertin, 125 mg/kg, i.p.) using 1XPBS (~30ml) and then 4% paraformaldehyde in 1XPBS (~50ml) via a Pharmacia pump (Cole-Parmer, Montreal, Canada). Brains were then dissected and post-fixed in 4% paraformaldehyde (overnight at 4°C), then cryoprotected in a 30% sucrose 1XPBS solution (48 hours at 4°C). Subsequently, the brains were frozen on dry ice until cryosectioning using a cryostat (Leica CM1950, Buffalo Grove, IL, USA). 20-μm-thick sections containing the PFC (AP 2.0–2.5 mm from Bregma) and then placed on the plus gold slides (Fischer scientific, #22035813) and stored at −80°C.

PFC sections were stained using primary antibodies against cell type markers specific for neurons (NeuN), astroglia (GFAP) and for GCaMP and RCaMP sensors (GFP and RFP respectively). Briefly, sections were washed three times in 1× PBS and then three times in PBS containing 0.3–0.5% Triton X-100 (PBS-T) and blocked for 30 min in 10% normal donkey serum (NDS) in PBS-T and then incubated with primary antibodies (See Supplementary Table1 for antibody combinations) in 1% NDS in PBS-T for 12 h at 4°C. After several washes in 1× PBS, sections were incubated with the corresponding secondary antibodies (See Supplementary Table1 for antibody combinations) for 2 h at room temperature and washed again in 1× PBS. Sections were coverslipped with Vectashield anti-fade mounting medium (Vector Laboratories) and kept at 4°C until imaging.

*Imaging and Quantification*

Images were acquired using a Olympus IX83 inverted microscope equipped with a spinning disk confocal unit and Hamamatsu Orca-Flash4.0 V2 digital CMOS camera. The images were captured at 20X magnification using SlideBook 6 imaging software (Intelligent Imaging Innovations, Denver, CO). Using the same exposure, 3-4 images per mouse were taken from the infected PFC regions (40 z stacks per image, 0.1 μm z-step size). Image’s origins (mouse ID and group) were blinded for downstream quantification. Using Fiji ImageJ software, images from each channel were superimposed and converted from slidebook to .tif format. Stained cells were quantified using QuPath (version 0.5.0) where all NeuN^+^ or GFAP^+^ cells within 332.8 x 332.8µm images were automatically traced and counted. The same process was used to quantify RFP^+^ and GFP^+^ infected cells and superimposed to NeuN^+^ and GFAP^+^ cell traces to identify co-labelled cells. Percentage of RFP^+^NeuN^+^/ total NeuN^+^ cells and GFAP⁺GFP⁺ / total GFAP⁺ were quantified to estimate infection rate of each virus.

**SUPPLEMENTARY RESULTS**

*Single-channel recordings*

To investigate the real-time modulation of cortical neuronal and/or astroglial Ca^2+^ activity during acute stress we performed *in vivo* fiber photometry using single-color (Figure S1). Following the experimental design illustrated in Figure S1A, in separate single-color cohorts (n = 8 each), the genetically encoded Ca^2+^ indicator GCaMP6f was expressed under either the neuron-specific human synapsin1 (Syn1) promoter or the astrocyte-specific GfaABC1D promoter using an AAV delivery system; Acute tail pinch robustly triggered PFC neuronal Ca²⁺ transients (Figure S1 B-E). To characterize the magnitude and kinetics of these tail pinch-evoked responses, we quantified the mean z-scored ΔF/F and the integrated Ca^2+^ signal (area under the curve, AUC) across three temporal windows relative to stimulus onset: pre-pinch (−5 to 0 s), pinch (0 to 3 s), and post-pinch (3 to 20 s). Repeated-measures one-way ANOVA revealed a significant main effect of time window on neuronal Ca^2+^ signal (z-scored ΔF/F: F_(2,14)_=27.67, *p*<0.0001; AUC: F_(2,14)_=37.91, *p*=0.0004) showing clear increases during pinch (z-scored ΔF/F: p < 0.01; AUC: p < 0.05, *Tukey’s post hoc*) and post-pinch (z-scored ΔF/F: p<0.01; AUC: *p*<0.0001) (Figure S1 D-E). Tail pinch-evoked responses were similar in astroglia with significant increase in mean z-scored ΔF/F (F_(2,14)_=8.994, *p*=0.01) and AUC (F_(2,14)_=17.80, *p*=0.003) during pinch (z-scored ΔF/F: *p*<0.01; AUC: *p*<0.05) and post-pinch (z-scored ΔF/F: *p*<0.01; AUC: *p*<0.01) (Figure S1 H-I). The viral cell-type specificity of each virus was confirmed by immunohistochemistry (Figure S1 J-K). These tail-pinch-evoked responses corroborate prior single-channel fiber photometry studies that recorded tail pinch-evoked activity in PFC neurons [4] or astrocytes [5]**.**

We next assessed the effects of a 30 min sustained immobilization stress (IS) on PFC neuronal or astroglial Ca²⁺ signals, recorded individually in the same animals as illustrated in Supplementary Figure 2A. In the Syn1-GCaMP cohort, IS triggered a significant increase in neuronal Ca^2+^ event frequency (Figure S2 B-D). Repeated-measures ANOVA revealed a main effect of time (F_(2,12)_=5.72, *p*<0.05) driven by a difference between pre-IS and IS conditions (p=0.01; Tukey’s *post-hoc* test, Figure S2 C). Similarly, in the GfaABC1D-GCaMP cohort, analysis showed main effect of time (F_(2,14)_=5.00, *p*<0.05) reflecting a significant increase in astroglia Ca^2+^ events during IS (*p*=0.02; Tukey’s *post-hoc* test, Figure S2 E-J). Event peak amplitude was similarly elevated during IS in neurons (F_(2,12)_=11.08, *p*<0.01, Supplementary Figure 2D) but did not reach to significance in astroglia.

*Additional behavioral data*

Hourly time spent in the shelter in phenotyper test for each week is presented in Supplementary Figure 3. Statistics obtained for weekly testing are reported in Supplementary Table 2. No change in water consumption was observed across weeks (Supplementary Figure 3G and Table S2). Locomotor activity was measured at week 5 and there was no difference between groups (no-UCMS:1803.6±120.26, UCMS: 1896.2±153.7).

*Weekly Ca^2+^ event changes*

Repeated measures ANOVA was used to assess the changes in event frequency and peak amplitude of neuronal (Supplementary Figure 4) or astroglial (Supplementary Figure 5) Ca^2+^ transients separately in no-UCMS control or UCMS group. Within-group and within-week statistics are reported in supplementary table 3

**SUPPLEMENTARY FIGURES**

**Supplementary Figure 1. Neuronal and Astroglial Ca^2+^ dynamics in response to tail pinch stimulus in mouse PFC. (A)** Schematic of experimental timeline from surgery to brain collection highlighting procedure up to the tail pinch. Panel A was created in  <https://BioRender.com>. **(B, F)** Average Z-scores ΔF/F of neuronal **(B)** and astroglial **(F)** Ca^2+^ activity from -5 to 20 sec with tail pinch stimuli applied from 0 to 3 sec. Solid lines indicate mean and shaded areas indicate s.e.m. **(C, G)** Heatmaps of Z-score ΔF/F of neuronal **(C)** and astroglial **(G).** Histograms illustrate quantified average Z-score ΔF/F and AUC of neuronal **(D, E)** and astroglial **(H, I)** Ca^2+^ signals calculated for each time period. Individual data points represent single subjects (circles for males, triangles for females). Data is represented as mean ± s.e.m. *p<0.05, **p<0.01, ***p<0.001, ****p<0.0001, Tukey *post-hoc* test. (**I-J)** Representative images of AAV-Syn1-GCAMP **(J)** and AAV-GFAP-GCaMP **(K)** infected areas labelled with GFAP (red), NeuN (blue) and GCaMP (green)**.** Both viruses were cell-specific and showed approximate infection rate of 80-85% under the cannula site.

**Supplementary Figure 2. Immobilization stress (IS) challenge increases neuronal and astroglial activity in PFC. (A)** Schematic of experimental timeline from surgery to brain collection highlighting IS. Panel A was created in  <https://BioRender.com>. **(B, E)** Heatmaps of Z-score ΔF/F (5 min-duration) of neuronal **(B)** and astroglial Ca^2^ **(E)** signals in the PFC obtained for each mouse. Chosen 5 min windows are the last 5min of pre-IS and the 5 first minutes of IS and Post-IS challenge. Histograms represent the Ca^2+^ event frequency and amplitude of PFC neurons **(C, D)** and astroglia **(F, G)** during each 30 min condition (pre-IS, IS and post-IS). Data is represented as mean ± s.e.m. Individual data points represent single mouse (circles for males, triangles for females). Tukey’s *post-hoc* test: *p<0.05, **p<0.01.

**Supplementary Figure 3. Light-challenge evoked residual avoidance in the PhenoTyper test and weekly water consumption.** Time spent in the shelter zone (sec) across the dark-phase recording (7 PM–7 AM) in the PhenoTyper test. The shaded bar with dashed outlines marks the 1 hr light challenge (11 PM–12 AM). **(A)** Shows the full cohort (all 12 mice) prior to randomization, plotted as a single curve. **(B-F)** Weekly time spent in the shelter zone for t no-UCMS and UCMS groups. **(G)** Weekly water consumption (ml) for both groups across Weeks 0–5. Data are presented as mean ± SEM. Tukey’s *post-hoc* test: **p* < 0.05, ***p* < 0.01, ****p* < 0.001; ^#^*p* < 0.1.

**Supplementary Figure 4. Weekly neuronal peak event frequency and peak amplitude across Pre, IS, and Post condition in response to IS challenge in no-UCMS control and UCMS group.** **(A)** Peak event frequency of neuronal ΔF/F Ca^2+^ signals across the Pre, IS (purple shaded region), and Post condition, shown separately for each week (weeks 0-5; columns) in no-UCMS mice (top row) and UCMS mice (bottom row). **(B)** Peak amplitude of neuronal events across the three conditions, for each week (column) and each group (row). Data is represented as mean ± s.e.m. Individual data points represent single mouse (circles: males, triangles: females). Tukey’s *post-hoc* test: **p* < 0.05, ***p* < 0.01, ****p* < 0.001.

**Supplementary Figure 5. Weekly astroglial peak event frequency and peak amplitude across Pre, IS, and Post condition in response to IS challenge in Control and UCMS cohort.** **A)** Peak event frequency of astroglial ΔF/F Ca^2+^ signals across the Pre, IS (purple shaded region), and Post condition, shown separately for each week (weeks 0-5; columns) in no-UCMS mice (top row) and UCMS mice (bottom row). **(B)** Peak amplitude of astroglial events across the three conditions, for each week (column) and each group (row). Data is represented as mean ± s.e.m. Individual data points represent single mouse (circles: males, triangles: females). Tukey’s *post-hoc* test: **p* < 0.05, ***p* < 0.01, ****p* < 0.001.

**Supplementary Figure 6. Repeated measure (*rmcorr*) correlation between neuron–astrocyte across weeks** Repeated-measures correlation (*rmcorr*) between per-minute neuronal and astroglial Ca²⁺ event frequencies; each panel shows the per-bin scatter of paired neuronal (x) and astroglial (y) event rates for individual-mouse data points with Pearson’s regression lines (shades of blue or pink) and the bold black line indicating the *rmcorr* common within-subject slope obtained for No UCMS animals **(A)** and UCMS animals **(B)**. Within **(A)** and **(B)** panels, each row corresponds to the three within-session conditions: pre-IS, IS and post-IS challenge. Columns correspond to weekly recording sessions (Week 0-5). The annotation box on the upper left side represents the r and p values and also summarized in Table 1 in main manuscript.
