## Supplementary Figures for "*In vivo* cortical neuron-astroglial functional coupling strengthens with acute stress but is impaired by chronic stress in mice"

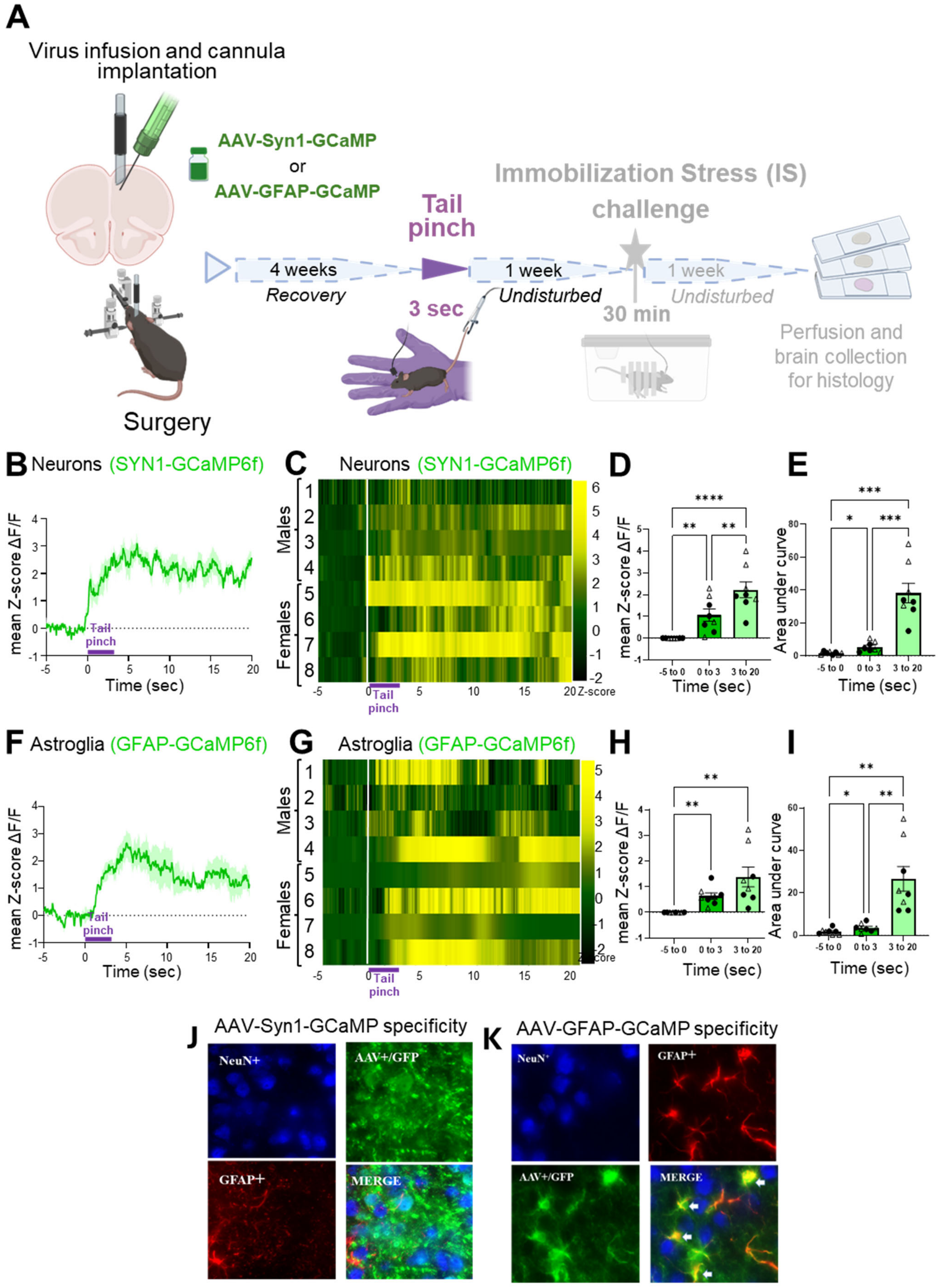


### Supplementary Figure 1

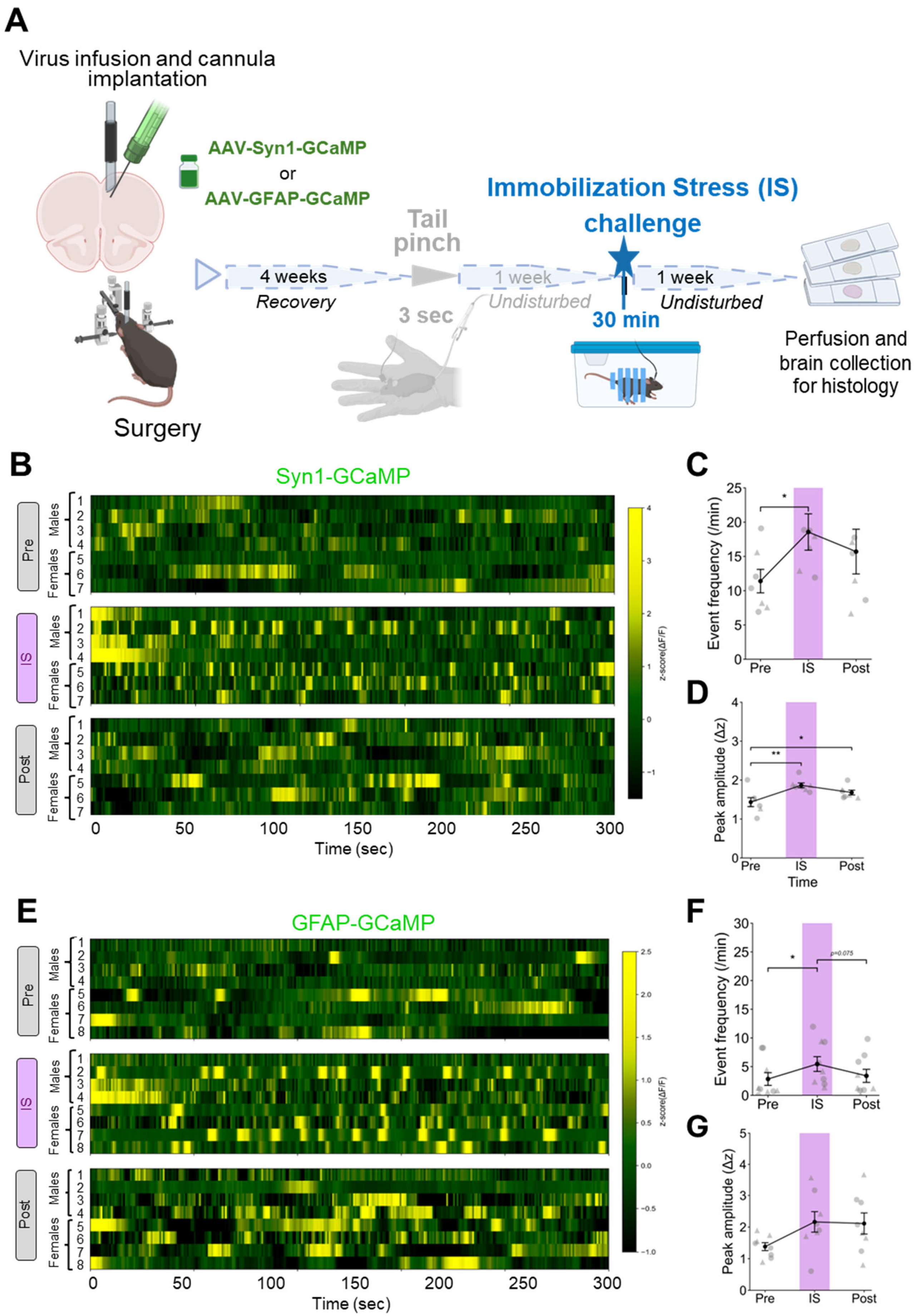


### Supplementary Figure 2

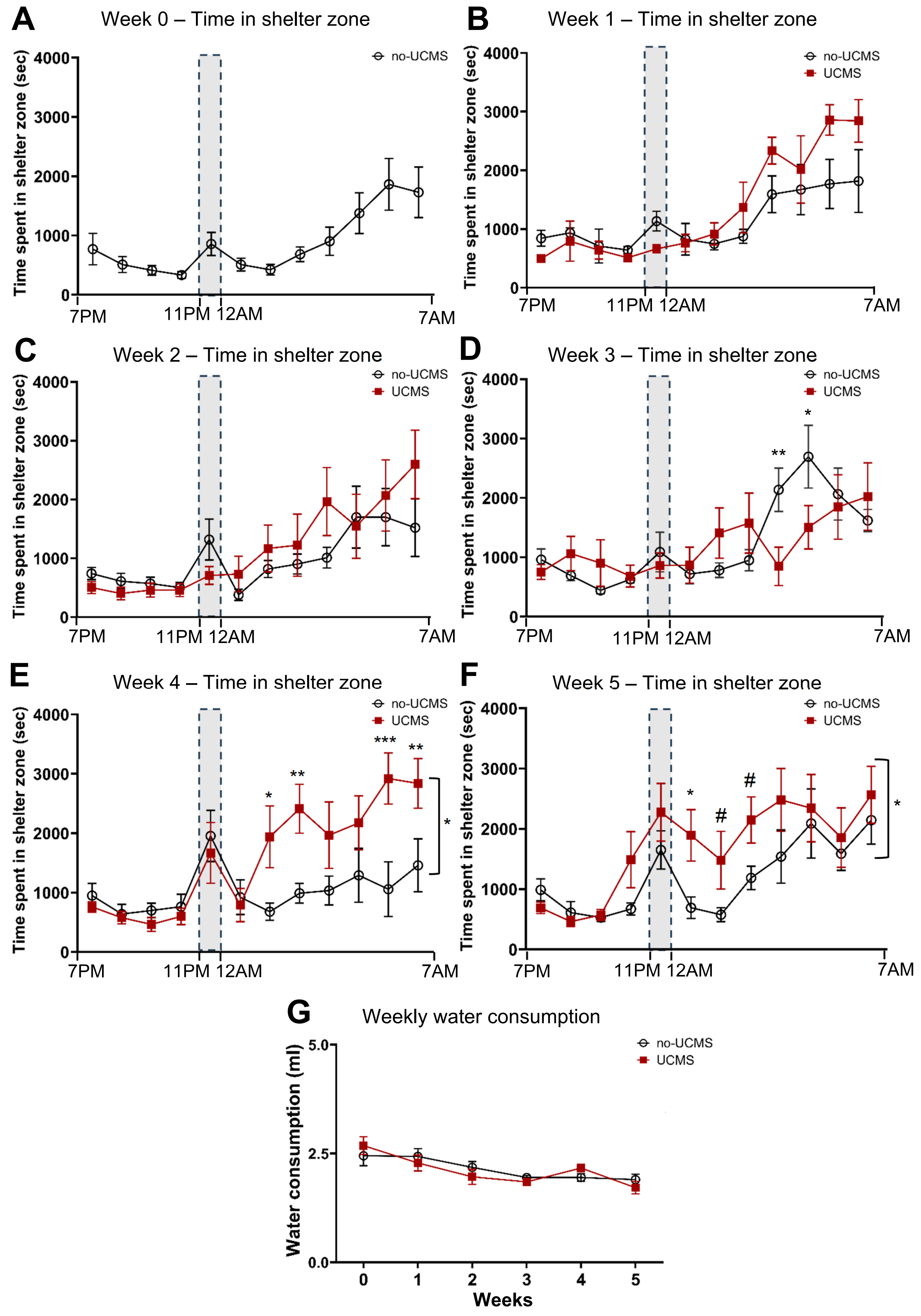


**Supplementary Figure 3**

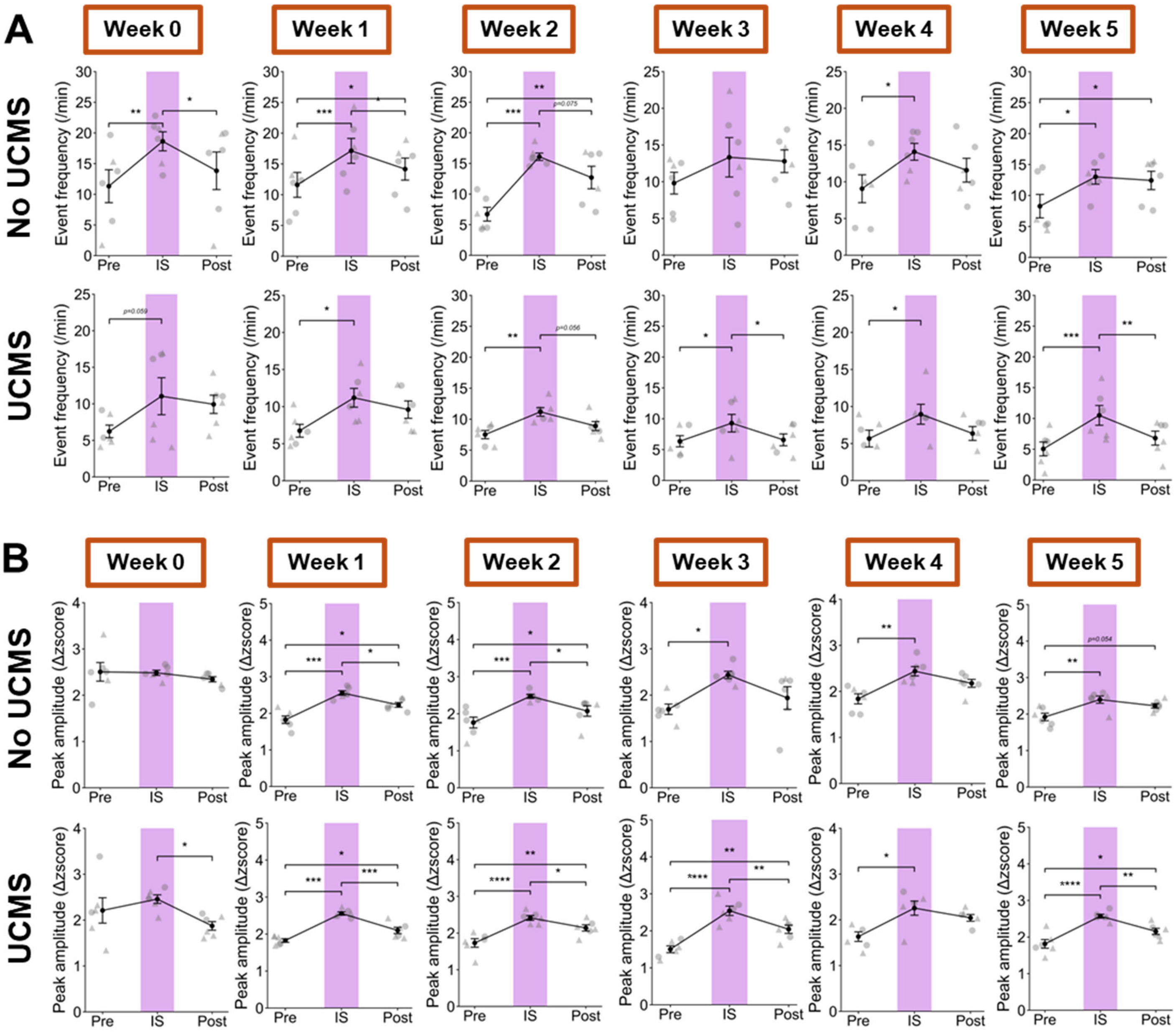


Supplementary Figure 4

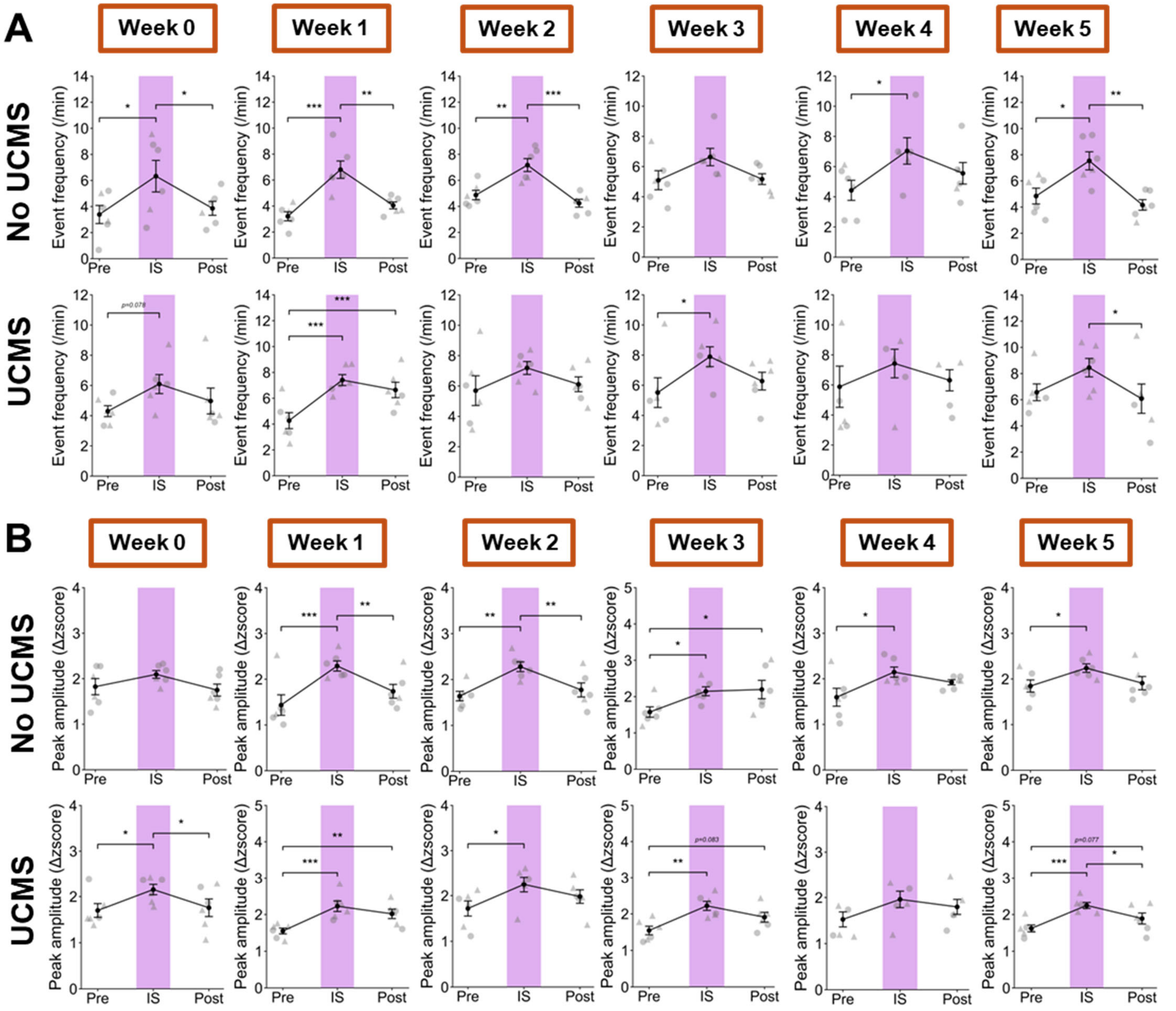


**Supplementary Figure 5**

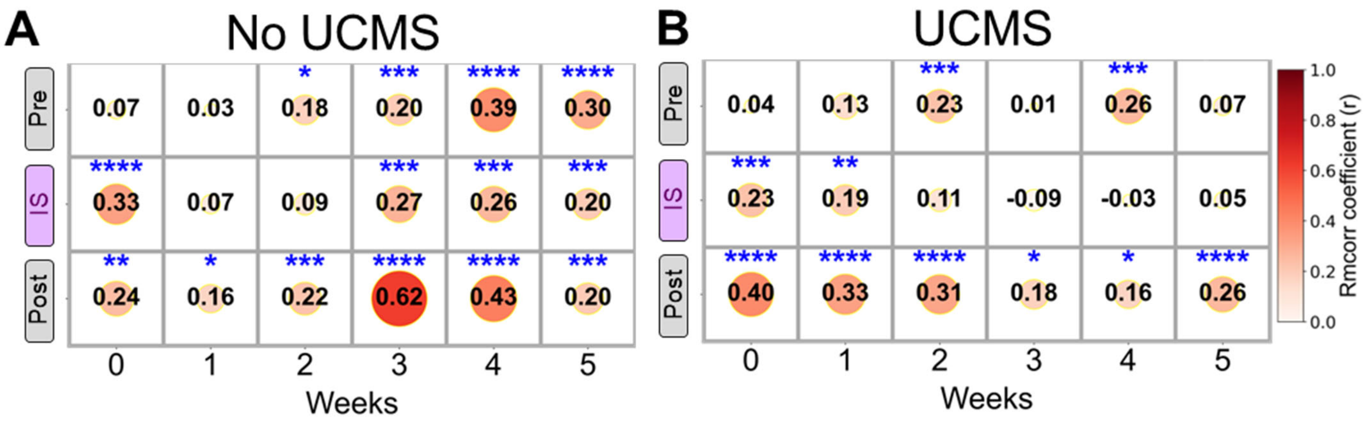
A
No UCMS
Pre
IS
Post
0.07
0.03
*
0.18
***
0.20
****
0.39
****
0.30
****
0.33
0.07
0.09
***
0.27
***
0.26
***
0.20
**
0.24
*
0.16
***
0.22
****
0.62
****
0.43
***
0.20
0
1
2
3
4
5
Weeks
B
UCMS
0.04
0.13
***
0.23
0.01
***
0.26
0.07
***
0.23
**
0.19
0.11
-0.09
-0.03
0.05
****
0.40
****
0.33
****
0.31
*
0.18
*
0.16
****
0.26
Weeks
Rmcorr coefficient (r)
1.0
0.8
0.6
0.4
0.2
0.0

**Table 1**
