## Supplementary Tables for "*In vivo* cortical neuron-astroglial functional coupling strengthens with acute stress but is impaired by chronic stress in mice"

| Purpose: specificity of virus | **Primary Antibody** | **Company - Catalog#** | **Secondary Antibody** | **Company - Catalog#** |
| --- | --- | --- | --- | --- |
| AAV-SYN1-GCaMP6f-WPRE-SV40 https://www.addgene.org/100837/ | Rb α GFAP (1:200) | Dako - Z0334 | Donkey α Rb AF 555 (1:200) | Invitrogen - A21572 |
|  | Gt α GFP (1:200) | Rockland - 600-101-215 | Donkey α Gt AF 488 (1:200) | Invitrogen - A11055 |
|  | GP α NeuN (1:500) | EMD Millipore - ABN90 | Donkey α GP AF 405 (1:200) | Biotium - 20376 |
| AAV5-Zac2.1 gfaABC1D-cyto-GCaMP6f https://www.addgene.org/52925/ | Rb α GFAP (1:200) | Dako - Z0334 | Donkey α Rb AF 555 (1:200) | Invitrogen - A21572 |
|  | Gt α GFP (1:200) | Rockland - 600-101-215 | Donkey α Gt AF 488 (1:200) | Invitrogen - A11055 |
|  | GP α NeuN (1:500) | EMD Millipore - ABN90 | Donkey α GP AF 405 (1:200) | Biotium - 20376 |
| AAV.Syn.NES.jRCaMP1a.WPRE.SV40 https://www.addgene.org/100848/ | Rb α GFAP (1:200) | Dako - Z0334 | Donkey α Rb AF 488 (1:200) | Invitrogen - A21206 |
|  | Chk α RFP (1:100) | Rockland - 600-401-379 | Donkey anti-Chk Rhodamine Red (1:200) | Jackson Immunoresearch - 703295155 |
|  | GP α NeuN (1:500) | EMD Millipore - ABN90 | Donkey α GP AF 405 (1:200) | Biotium - 20376 |
|  | Rb = Rabbit, Chk = Chicken, GT= Goat, Glial Fibrillary Acidic Protein (GFAP), Green Fluorescent Protein (GFP), Red Fluorescent Protein (RFP) | | | |

**Table 1: List of Antibodies**

**Table 2: Statistical analysis of the supplementary data**

| Experiment | Week of Testing | Statistics (F,p) | Group differences: Tukey's post-hoc analysis |
| --- | --- | --- | --- |
| Phenotyper test | Week0 | Main effect of Time: F(11,110) =7.54, p=0.0004 | No group differences |
|  | Week1 | UCMS: F(1,10) =1.74, p = ns Main effect of Time: F(11,110) =12.19, p<0.0001 UCMSxTime interaction: F(11,110) =1.84, p = ns | No group differences |
|  | Week2 | UCMS: F(1,10) =0.50, p = ns Main effect of Time: F(11,110) =6.32, p=0.0008 UCMSxTime interaction: F(11,110) =1.205, p = ns | No group differences |
|  | Week3 | UCMS: F(1,10) =0.04, p = ns Main effect of time: F(11,110) =5.23, p<0.0001 UCMSxTime interaction: F(11,110) =2.05, p=0.029 | 3am: No-UCMS vs UCMS**, 4am No-UCMS vs UCMS* |
|  | Week4 | UCMS: (F(1,10) =5.48, p<0.05) Main effect of Time: F(11,110) =7.65, p<0.0001 UCMSxTime interaction: F(11,110) =3.154, p=0.0006 | 1am: No-UCMS vs UCMS*, 2am No-UCMS vs UCMS**, 5am No-UCMS vs UCMS***, 6am No-UCMS vs UCMS**, 7am No-UCMS vs UCMS* |
|  | Week5 | UCMS: F(1,10) =9.65, p<0.05 Main effect of Time: F(11,110) =6.38, p=0.011 UCMSxTime interaction: F(11,110) = 0.91, p=ns | 12am: No-UCMS vs UCMS*, 1am: No-UCMS vs UCMS#, 2am No-UCMS vs UCMS# |
| Water consumption test |  | Time*UCMS interaction: (F(5,50) =0.96 p=449) | No group differences |
|  |  | #p < 0.1, *p < 0.05, **p < 0.01, ***p < 0.001, |  |

**Table 3: Statistical analysis of the supplementary neuronal and astroglial Ca2+ activity data**

| Experiment | Week of Testing | Groups | Parameter | Statistics (F,p) | Condition differences: Tukey's post-hoc analysis |
| --- | --- | --- | --- | --- | --- |
| Weekly neuronal peak analysis | Week0 | no-UCMS | Peak frequency | Condition: (F(2,10) =13.37, p=0.0015) | Pre-IS vs IS**, IS vs Post-IS** |
|  |  |  | Amplitude | Condition : (F(2,10) =0.45, p=0.646) | No difference |
|  |  | UCMS | Peak frequency | Condition : (F(2,10) =3.10, p=0.08) | Pre-IS vs IS(0.058) |
|  |  |  | Amplitude | Condition : (F(2,10) =6.18, p=0.017) | IS vs Post-IS* |
|  | Week1 | no-UCMS | Peak frequency | Condition : (F(2,10) =19.84, p=0.0003) | Pre-IS vs IS***, IS vs Post-IS**, Pre vs Post-IS* |
|  |  |  | Amplitude | Condition : (F(2,10) =19.12, p=0.0004) | Pre-IS vs IS***, IS vs Post-IS*, Pre vs Post-IS* |
|  |  | UCMS | Peak frequency | Condition : (F(2,10) =4.407, p=0.04) | Pre-IS vs IS* |
|  |  |  | Amplitude | Condition : (F(2,10) =41.81, p<0.0001) | Pre-IS vs IS***, IS vs Post-IS***, Pre vs Post-IS* |
|  | Week2 | no-UCMS | Peak frequency | Condition : (F(2,10) =24.34, p=0.0001) | Pre-IS vs IS***, Pre-IS vs Post-IS** |
|  |  |  | Amplitude | Condition : (F(2,10) =20.30, p=0.0003) | Pre-IS vs IS***, IS vs Post-IS*, Pre vs Post-IS* |
|  |  | UCMS | Peak frequency | Condition : (F(2,10) =9.396, p=0.005) | Pre-IS vs IS**, IS vs Post-IS(0.056) |
|  |  |  | Amplitude | Condition : (F(2,10) =35.24, p<0.0001) | Pre-IS vs IS****, IS vs Post-IS**, Pre vs Post-IS* |
|  | Week3 | no-UCMS | Peak frequency | Condition : (F(2,10) =3.35, p=0.078) | No difference |
|  |  |  | Amplitude | Condition : (F(2,10) =5.91, p=0.02) | Pre-IS vs IS* |
|  |  | UCMS | Peak frequency | Condition : (F(2,10) =6.029, p=0.01) | Pre-IS vs IS*, Pre-IS vs Post-IS* |
|  |  |  | Amplitude | Condition : (F(2,10) =35.78, p<0.0001) | Pre-IS vs IS****, IS vs Post-IS**, Pre vs Post-IS** |
|  | Week4 | no-UCMS | Peak frequency | Condition : (F(2,10) =4.499, p=0.04) | Pre-IS vs IS* |
|  |  |  | Amplitude | Condition : (F(2,10) =8.09, p=0.008) | Pre-IS vs IS** |
|  |  | UCMS | Peak frequency | Condition : (F(2,10) =4.74, p=0.036) | Pre-IS vs IS* |
|  |  |  | Amplitude | Condition : (F(2,10) =6.67, p=0.014) | Pre-IS vs IS* |
|  | Week5 | no-UCMS | Peak frequency | Condition : (F(2,10) =6.024, p=0.04) | Pre-IS vs IS*, Pre-IS vs Post-IS* |
|  |  |  | Amplitude | Condition : (F(2,10) =9.01 p=0.005) | Pre-IS vs IS**, Pre vs Post-IS(0.054) |
|  |  | UCMS | Peak frequency | Condition : (F(2,10) =22.45, p=0.0009) | Pre-IS vs IS***, IS vs Post-IS** |
|  |  |  | Amplitude | Condition : (F(2,10) =30.37, p<0.0001) | Pre-IS vs IS****, IS vs Post-IS**, Pre vs Post-IS* |
| Weekly astroglial peak analysis | Week0 | no-UCMS | Peak frequency | Condition : (F(2,10) =7.074, p=0.012) | Pre-IS vs IS*, IS vs Post-IS* |
|  |  |  | Amplitude | Condition : (F(2,10) =1.88, p=0.201) | No difference |
|  |  | UCMS | Peak frequency | Condition : (F(2,10) =3.83, p=0.058) | Pre-IS vs IS(0.058) |
|  |  |  | Amplitude | Condition : (F(2,10) =6.91, p=0.013) | IS vs Post-IS* |
|  | Week1 | no-UCMS | Peak frequency | Condition : (F(2,10) =17.10, p=0.0006) | Pre-IS vs IS***, IS vs Post-IS*** |
|  |  |  | Amplitude | Condition : (F(2,10) =18.52, p=0.0004) | Pre-IS vs IS***, IS vs Post-IS** |
|  |  | UCMS | Peak frequency | Condition : (F(2,10) =29.69, p<0.0001) | Pre-IS vs IS***, Pre vs Post-IS*** |
|  |  |  | Amplitude | Condition : (F(2,10) =19.39, p=0.0004) | Pre-IS vs IS***, IS vs Post-IS** |
|  | Week2 | no-UCMS | Peak frequency | Condition : (F(2,10) =15.94, p=0.0008) | Pre-IS vs IS**, IS vs Post-IS*** |
|  |  |  | Amplitude | Condition : (F(2,10) =14.95, p=0.001) | Pre-IS vs IS**, IS vs Post-IS** |
|  |  | UCMS | Peak frequency | Condition : (F(2,10) =1.37, p=0.31) | No difference |
|  |  |  | Amplitude | Condition : (F(2,10) =4.09, p=0.05) | Pre-IS vs IS* |
|  | Week3 | no-UCMS | Peak frequency | Condition : (F(2,10) =2.68, p=0.116) | No difference |
|  |  |  | Amplitude | Condition : (F(2,10) =7.18, p=0.001) | Pre-IS vs IS*, Pre vs Post-IS* |
|  |  | UCMS | Peak frequency | Condition : (F(2,10) =4.05, p=0.05) | Pre-IS vs IS* |
|  |  |  | Amplitude | Condition : (F(2,10) =10.16, p=0.004) | Pre-IS vs IS**, Pre vs Post-IS(0.08) |
|  | Week4 | no-UCMS | Peak frequency | Condition : (F(2,10) =3.82, p=0.058) | Pre-IS vs IS* |
|  |  |  | Amplitude | Condition : (F(2,10) =4.42, p=0.04) | Pre-IS vs IS* |
|  |  | UCMS | Peak frequency | Condition : (F(2,10) =1.046, p=0.386) | No difference |
|  |  |  | Amplitude | Condition : (F(2,10) =1.58, p=0.252) | No difference |
|  | Week5 | no-UCMS | Peak frequency | Condition : (F(2,10) =10.62, p=0.007) | Pre-IS vs IS*, IS vs Post-IS** |
|  |  |  | Amplitude | Condition : (F(2,10) =4.65, p=0.037) | Pre-IS vs IS* |
|  |  | UCMS | Peak frequency | Condition : (F(2,10) =4.79, p=0.054) | IS vs Post-IS* |
|  |  |  | Amplitude | Condition : (F(2,10) =16.74, p=0.0006) | Pre-IS vs IS***, IS vs Post-IS*, Pre vs Post-IS(0.07) |
|  |  | UCMS: Unpredictable Chronic Mild Stress, IS: Immobilization stress, *p < 0.05, **p < 0.01, ***p < 0.001, ****p < 0.0001 | | | |
